# Titin cleavage impairs cardiac mechanical connectivity to drive diastolic failure and fibrosis

**DOI:** 10.1101/2025.04.21.649485

**Authors:** Johanna K. Freundt, Paulina Hartmann, Christine M. Loescher, Andreas Unger, Franziska Koser, Annika J. Klotz, Lydia Wachsmuth, Susanne Hille, Michaela M. Door, Anne Helfen, Richard Holtmeier, Cornelius Faber, Jonathan A. Kirk, Verena Hoerr, Oliver J. Müller, Wolfgang A. Linke

## Abstract

Titin, the largest human protein, serves as the elastic backbone of sarcomeres in muscle cells and imparts passive stiffness to cardiomyocytes. While proteolytic cleavage of elastic titin has been linked to cardiovascular disease, the isolated effects of titin stiffness loss in the living heart remain poorly understood. Here, we developed a knock-in mouse model allowing for the selective cleavage of cardiac titin springs in vivo. Using a time-resolved approach combining MRI, echocardiography, immunofluorescence, and molecular profiling, we demonstrate that titin cleavage does not dilate the heart but induces concentric remodeling, impaired diastolic function, and low-output heart failure. Mechanistically, compromised titin-based restoring forces trigger internal mechanical imbalance and dysconnectivity, which activate fibroblasts and promote extracellular matrix remodeling, revealing titin’s essential role in maintaining cardiac mechanical homeostasis.

## Main

The giant protein titin provides structural integrity to the sarcomere, the contractile unit of striated muscle (*1–4*), and underlies its viscoelastic properties (*5, 6*). Spanning half a sarcomere from the Z-disk to the M-band, titin governs the passive stiffness of cardiomyocytes (*7–9*) and modulates contraction (*5*). Titin alterations are implicated in cardiomyopathy and heart failure, with gene truncations in *Ttn* accounting for many inherited cases (*10*). Shifts between its principal isoforms, N2BA and N2B (*11, 12*), together with post-translational modifications (*13*), have been linked to various heart failure syndromes. Moreover, proteolytic cleavage within titin’s elastic I-band has been associated with atrial fibrillation (*14*) and myocardial injury from chemotherapies (*15, 16*) or ischemia (*17, 18*). However, the direct impact of titin stiffness loss – such as via I-band cleavage – remains speculative because its study requires isolated cleavage of titin springs in vivo without disrupting sarcomere scaffolding, as seen in titin-deletion models (*2, 19*).

Another unresolved question concerns the mechanical interplay between cardiomyocytes and the extracellular matrix (ECM) and whether titin stiffness influences this interaction. Although myocardial damage frequently leads to fibrosis (*20*) and affects structural remodeling during heart failure progression (*21, 22*), it remains unclear how changes in cardiomyocyte force generation directly affect ECM remodeling (*23, 24*). Activated fibroblasts (myofibroblasts), which secrete collagen – the major stiffening ECM component – respond acutely to mechanical cues such as substrate stiffness or stretch (*25*), regulating ECM stiffness through homeostatic feedback loops (*26*). Furthermore, diminished cardiomyocyte contractility may be sensed by matrix components, triggering adaptive ECM remodeling to restore optimal mechanical conditions (*23*). Whether alterations in titin-based passive stiffness can initiate such a response remains an open question. To address this, we selectively cleaved cardiac titin springs in vivo to assess their impact on ventricular mechanics and potential cross-talk with fibroblasts and ECM remodeling.

### Specific cleavage of elastic titin in the beating mouse heart

We hypothesized that selective cleavage of elastic titin would recapitulate aspects of titin-related cardiomyopathy and elucidate the role of titin stiffness in maintaining cardiac mechanical balance. To test this, we employed the titin cleavage (TC) mouse model (*27*), inserting a tobacco etch virus protease (TEV) recognition site into titin’s I-band region between immunoglobulin-like domains 86 and 87 (**Fig. 1A**). Myocardial-specific cleavage was induced via AAV9-mediated TEV expression under a cardiac troponin-T promoter, delivered by tail-vein injections into homozygous (Hom) or heterozygous (Het) TC mice, with AAV9-GFP serving as a control. Cardiac function was monitored by magnetic resonance imaging (MRI) at day (D)6 and D13 post injection and by transthoracic echocardiography (TTE) at D0, D3, D6, D10, and D13 (**Fig. 1B**). Voluntary running activity was recorded throughout, and tissue samples were mainly collected at D6 and D13.

**Figure 1.**
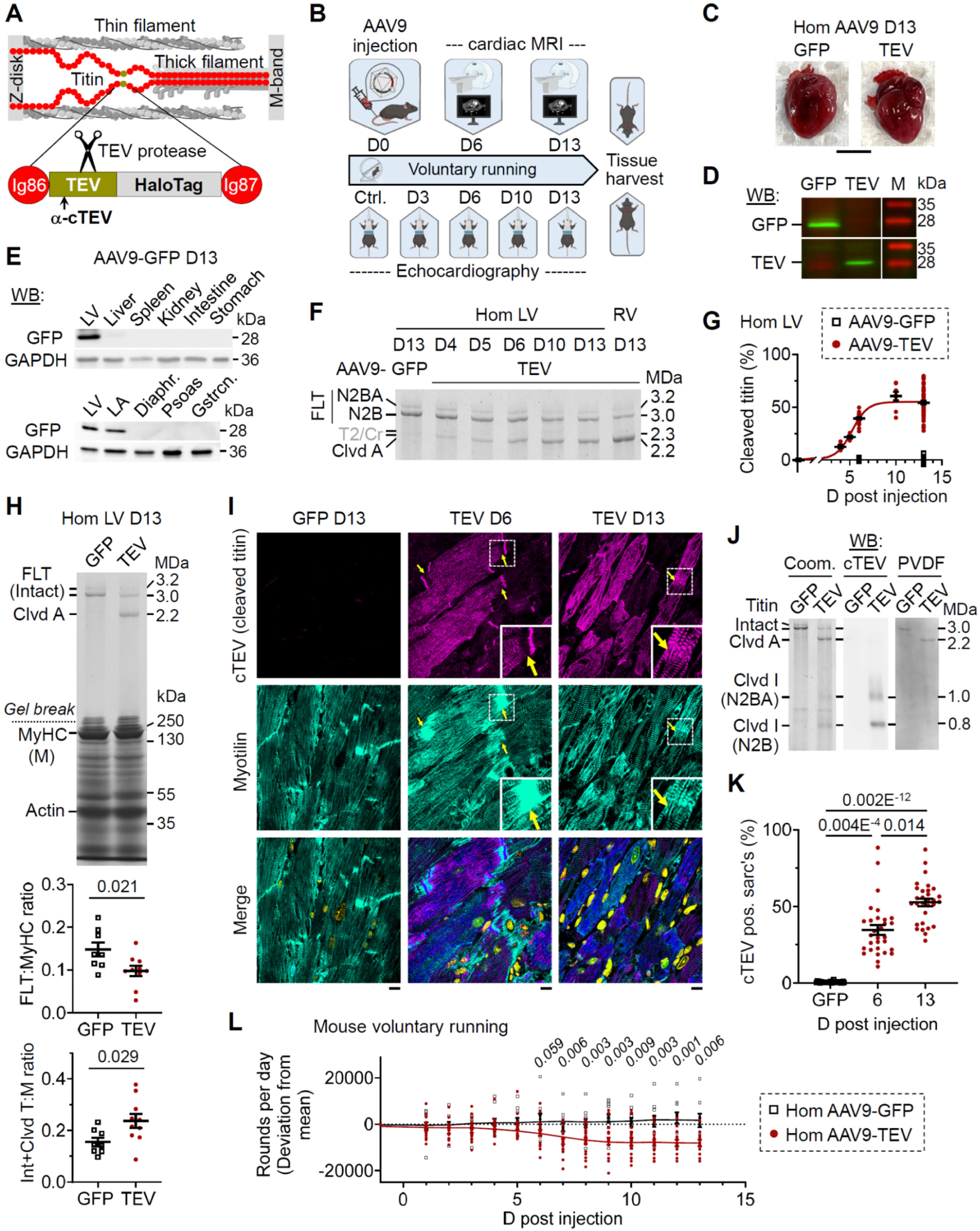
In vivo titin cleavage model. **(A)** Schematic of a TC mouse half-sarcomere showing the TEV protease recognition site in titin’s I-band; arrow indicates the cleaved titin recognized by α-cTEV. **(B)** Experimental timeline: AAV9-TEV or AAV9-GFP (control) plasmids were injected into homozygous (Hom) or heterozygous (Het) TC mice at D0; voluntary running was recorded; cardiac MRI was performed at D6 and D13, echocardiography at D0, D3, D6, D10, and D13; tissues were collected mainly at D6 and D13. **(C)** Explanted heart images from AAV9-GFP and AAV9-TEV Hom TC mice at D13; scale bar, 5 mm. **(D)** Western blot (WB) confirming GFP and TEV overexpression in heart tissues at D13 (Hom TC). **(E)** WB analysis of GFP expression in various organs and muscles from D13 AAV9-GFP mice; GAPDH is the loading control (LV, left ventricle; LA, left atrium; Diaphr., m. diaphragm; Gstrcn., m. gastrocnemius). **(F)** Coomassie-stained titin gels showing the time-dependent increase in titin cleavage in LV and right ventricle (RV) of Hom TC mice following AAV9-TEV injection versus AAV9-GFP; N2BA and N2B denote intact (full-length) titin isoforms, Clvd A indicates cleaved titin (mostly A-band). T2/Cr refers to titin proteolytic fragment and Cronos. **G)** Time course of titin cleavage: AAV9-GFP mice: N=7 (n=7) at D0, N=6 (n=6) at D6, N=17 (n=28) at D13; AAV9-TEV mice: N=1 (n=6) at D4-D5, N=6 (n=23) at D6, N=2 (n=8) at D10, and n=17 (n=66) at D13. **(H)** Coomassie-stained protein gel (*upper*: titin; *lower*: standard) of LV proteins at D13 post injection with graphs of intact titin/MyHC (*middle*) and total titin/MyHC (*bottom*); N=8 (GFP), N=10 (TEV); significance by unpaired two-tailed t-test. **(I)** Confocal images of LV tissue: *left*, D13 AAV9-GFP; *center*, D6; *right*, D13 post AAV9-TEV. *Top*: α-cTEV staining (Cy3); *middle*: α-Myotilin staining (AlexaFluor 488); *bottom*: merge with DAPI. Arrows indicate lost contact between titin-cleaved sarcomeres and intercalated disks (“streaming”). Scale bars, 10 µm; insets show magnified ROIs. **(J)** Confirmation of in vivo titin cleavage in Hom TC LV at D13 post AAV9-TEV versus GFP: *left*, Coomassie-stained titin gel (1.8% PA); *middle*, WB with α-cTEV; *right*, PVDF staining as loading control. **(K)** Quantitation of cleaved versus non-cleaved sarcomeres based on cTEV signals normalized to Myotilin (semi-automated Nikon NIS software). n=30 images (∼13,000– 18,000 sarcomeres/image); N=2 (GFP) and N=3 (TEV); significance by Kruskal-Wallis test with Dunn’s multiple comparisons. **(L)** Voluntary running activity: laps/day (deviation from start) in Hom TC mice post AAV9-GFP or AAV9-TEV injection; N=7-12 (GFP) and N=13-20 (TEV); significance by unpaired two-tailed t-test. All average data are presented as mean ± SEM.

At D13, hearts from AAV9-GFP and AAV9-TEV-injected Hom TC mice displayed similar external morphology (**Fig. 1C**) and robust expression of GFP and TEV, respectively (**Fig. 1D**). Western blots (WBs) using tissue from GFP-injected mice confirmed cardiac-restricted expression: GFP was evident in all heart chambers yet nearly absent in skeletal muscle, spleen, and even liver (**Fig. 1E**, **Fig. S1A**). In TEV-injected Hom TC left ventricles, titin cleavage, assessed on loose protein gels, increased from ∼12% at D4 to ∼40% at D6, plateauing at ∼55-60% by D10-D13 (**Fig. 1F-G**); titin was also cleaved in the right ventricle (RV). No cleavage was observed in GFP-injected controls. Both major titin isoforms (N2BA and N2B) were cleaved similarly, with isoform ratios nearly unchanged (**Fig. S1B**). Notably, cleaved titin levels exceeded the loss of intact titin, resulting in a net increase in total titin protein (**Fig. 1H**). TEV-cleaved titin appeared on WBs as an N-terminal fragment (Cleaved-I) and a C-terminal fragment (Cleaved-A) (**Fig. S1C-D**); Cleaved-A appeared as a single band, while Cleaved-I split into two signals corresponding to N2BA and N2B (**Fig. S1D**). The alternative titin isoform Cronos, abundantly expressed in the atria and fast skeletal muscles (*2*), was cleaved at its extreme N-terminus, confirming cleavage in the atria but not in skeletal muscles (**Fig. S1E**). Together, these data demonstrate the successful and specific in vivo cleavage of cardiac titin.

We next examined the spatial pattern of titin cleavage within cardiac tissue using immunofluorescence on left ventricular (LV) sections from Hom TC mice. An antibody that recognizes the TEV-cleaved sequence (α-cTEV) revealed a mosaic pattern of titin-cleaved and un-cleaved cardiomyocytes (**Fig. 1I**). On WBs, α-cTEV specifically detected the two Cleaved-I bands (**Fig. 1J**). Cleaved titin was localized within the sarcomere and at the intercalated disk (ICD), where co-staining with the Z-disk/ICD marker Myotilin revealed “streaming” at ICDs (**Fig. 1I**, arrows). Quantitation showed ∼35% cleaved sarcomeres at D6 and ∼53% at D13, consistent with WB data, while GFP-injected hearts were negative (**Fig. 1K**). Furthermore, TEV-injected LV exhibited a reduction in sarcomere density by 8.9% at D6 and 26.4% at D13 compared to controls (**Fig. S1F-G**). Functionally, TEV-injected Hom TC mice demonstrated reduced voluntary wheel running activity starting at D6-D7 (**Fig. 1L**), prompting further detailed cardiac functional analyses.

### Titin-cleaved hearts show concentric remodeling and diastolic failure

Time-resolved noninvasive imaging revealed that selective titin cleavage in TEV-injected versus GFP-injected Hom TC mice induced marked concentric remodeling – an unexpected finding given the proposed role of titin stiffness in limiting chamber distensibility (*5, 6*), yet consistent with its predicted function in supporting elastic recoil (*28, 29*). Short-axis cMRI scans (**Fig. 2A**) demonstrated pronounced reductions in both end-diastolic and end-systolic LV volumes at D6 and D13; diastolic (but not systolic) volume was even lower at D13 (**Fig. 2B**). The interventricular septum remained unchanged at end-diastole but was thicker at end-systole at D6 (**Fig. 2C**), while end-diastolic and end-systolic outer heart diameters were modestly decreased (**Fig. 2D**). The Green-Lagrange strain exhibited a sharp increase at D6 that diminished by D13 (**Fig. 2E**). Under anesthesia, heart rate (∼500 beats per minute) remained steady, yet stroke volume and cardiac output were markedly reduced at D6, with further declines by D13 (**Fig. 2F-G**). Notably, minor sex-dependent differences emerged, as females exhibited slightly greater reductions in LV end-diastolic volume, stroke volume, and cardiac output at D6 (**Fig. S2**). Cinematic long-axis cMRI scans (**Movies S1-S3**) showed minimal post-systolic expansion in titin-cleaved hearts, impairing LV inflow across the mitral valve. Moreover, pleural effusions frequently appeared at D13 (**Fig. 2A**), indicating severe backward failure.

**Figure 2.**
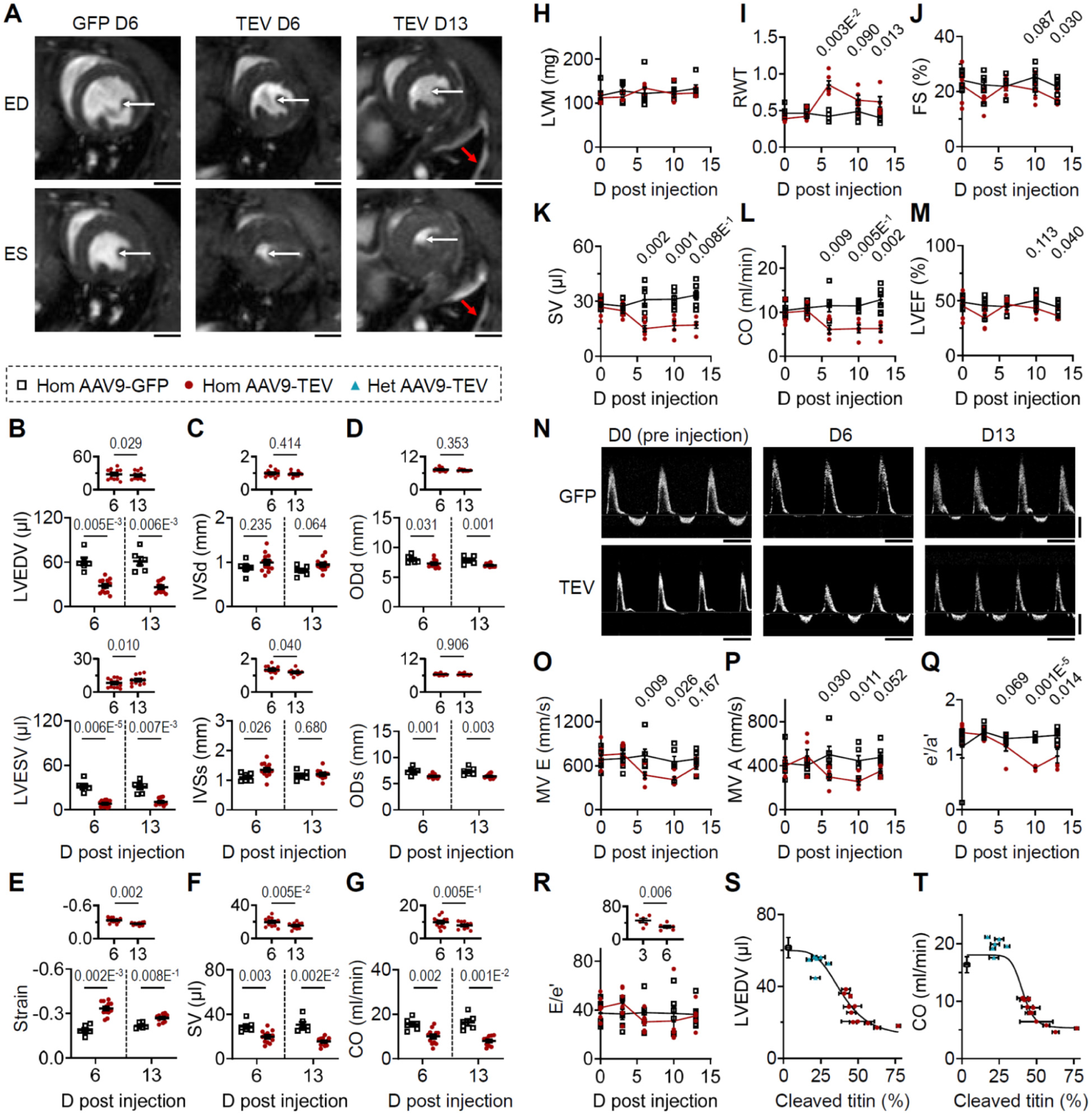
Cardiac function in AAV9-TEV titin-cleaved and AAV9-GFP control TC mice. **(A)** Representative short-axis cMRI images (two-chamber view) at end-diastole (ED) and end-systole (ES) at D6 and D13 post-AAV9 injection versus AAV9-GFP controls. Left ventricular (LV) volume (white arrows), interventricular septum (IVS) width, and outer heart diameter (OD) were quantified. Note pleural effusion (red arrows) at D13 post-TEV injection. Scale bar, 2 mm. **(B-G)** cMRI-derived parameters: (B) LVEDV and LVESV; (C) IVS (diastolic and systolic); (D) OD (diastolic and systolic); (E) Green-Lagrange strain; (F) stroke volume (SV); (G) cardiac output (CO). Small graphs show direct comparison of AAV9-TEV mice at D6 and D13. **(H-M)** M-mode TTE parameters: (H) LV mass (LWM); (I) relative wall thickness (RWT); (J) fractional shortening (FS); (K) SV; (L) CO; and (M) LV ejection fraction (LVEF). **(N)** Representative PW Doppler measurements of mitral valve (MV) flow at D0, D6, and D13; scale bars: 500 mm/s (vertical), 0.1 s (horizontal). **(O-P)** PW Doppler: (O) MV E wave velocity (early filling) and (P) MV A wave velocity (late filling). **(Q-R)** Tissue Doppler: (Q) e′/a′ ratio and (R) E/e′ ratio (small graph: direct comparison at D3 and D6 in AAV9-TEV mice). **(S-T)** Correlation of titin cleavage with core echocardiographic parameters: (S) LVEDV and (T) CO across AAV9-injected Hom and Het TC mice; lines represent sigmoidal fits to the mean data points. All data are presented as mean ± SEM. cMRI: N=6 (GFP), N=13 (D6 TEV), N=11 (D13 TEV); TTE: N=5–6 mice/group. Statistical significance was determined by unpaired two-tailed or paired t-tests (small graphs).

M-mode echocardiography confirmed the marked reduction in LV cavity size beginning at D6, accompanied by transient increases in septal and posterior wall thickness that partially recovered at later time points (**Fig. S3A-D**). LV mass remained unchanged (**Fig. 2H**), while relative wall thickness increased from D6 onward (**Fig. 2I**). Whereas fractional shortening declined only at later times (**Fig. 2J**), stroke volume and cardiac output were substantially reduced from D6 to D13 (**Fig. 2K-L**), with heart rate stable (**Fig. S3E**) and LVEF preserved except for a slight decline at D13 (**Fig. 2M**). Altogether, these findings indicate that titin cleavage initially preserves systolic function while impairing diastolic performance, followed by partial compensation in some parameters but progressive deterioration of cardiac contractility over time.

Doppler imaging (**Fig. 2N**) revealed diminished mitral valve flow, with both early (MV E, **Fig. 2O**) and late (MV A, **Fig. 2P**) filling velocities reduced from D6 to D10, followed by partial recovery, and a preserved E/A ratio (**Fig. S3F**). The pressure half time was prolonged and its acceleration slowed at D6, returning to baseline by D13 (**Fig. S3G-H**). Although early diastolic (e’) and atrial “kick” (a’) waves exhibited only minor changes (**Fig. S3I-J**), the e’/a’ ratio was noticeably lowered (**Fig. 2Q**), while isovolumic relaxation time remained largely unaltered (**Fig. S3K**). Doppler assessment also showed a transient decrease in the E/e’ ratio, a measure of LV filling pressure, at D6 (**Fig. 2R**), alongside reduced aortic flow velocity and lower aortic valve pressure gradients (**Fig. S3L-P**), collectively indicating low-output failure due to restricted diastolic filling.

To determine the threshold of titin cleavage required to alter key morphological and functional parameters, we also analyzed Het TC mice, in which TEV expression cleaved ∼27% of titin at D13 (**Fig. S4A**). Most biochemical and functional measures remained unchanged between TEV- and GFP-injected Het mice, although trends toward increased total titin expression (**Fig. S4B**) and reduced voluntary running activity could be observed (**Fig. S4C**). Notably, the Z-disk fracture score (*30*) – a sensitive marker of subcellular disruption – was markedly elevated in TEV-injected Het hearts (**Fig. S4D**), whereas no echocardiographic parameters were affected (**Fig. S4E-K**). Plotting parameter changes against the percentage of titin cleavage (in Het and Hom mice) revealed that LV end-diastolic volume and cardiac output were preserved until cleavage exceeded ∼30% (**Fig. 2S-T**), demonstrating the heart’s remarkable compensatory capacity.

### Reduced passive stiffness and restoring forces of titin-cleaved cardiomyocytes

Intrigued by the functional phenotype of titin-cleaved hearts, we isolated single cardiomyocytes from Het TC hearts (50% cleavable titins) for ex vivo mechanical analysis. Cells were permeabilized, suspended between a force transducer and micromotor, and incubated with TEV for 20-30 minutes, which cleaved roughly half the titin springs in a controlled manner (**Fig. S5A-B**). This intervention reduced stretch-dependent passive force by ∼23% (**Fig. S5C**), while Ca²⁺-activated force (pCa 5) remained unchanged compared to TEV-treated wildtype cardiomyocytes (**Fig. S5D**).

We next assessed the ability of permeabilized, lightly adherent Hom TC cardiomyocytes to recover sarcomere length (SL) after contraction. Cells activated at pCa 6 and then rapidly relaxed with EGTA (**Fig. 3A**) displayed full SL recovery over two cycles in un-cleaved cells (**Fig. 3B-C**); however, titin cleavage after the first cycle significantly impaired SL restoration during the second (p<0.005; **Fig. 3D-E**). These results indicate that titin cleavage diminishes restoring forces, impairing elastic recoil and thereby compromising ventricular filling.

**Figure 3.**
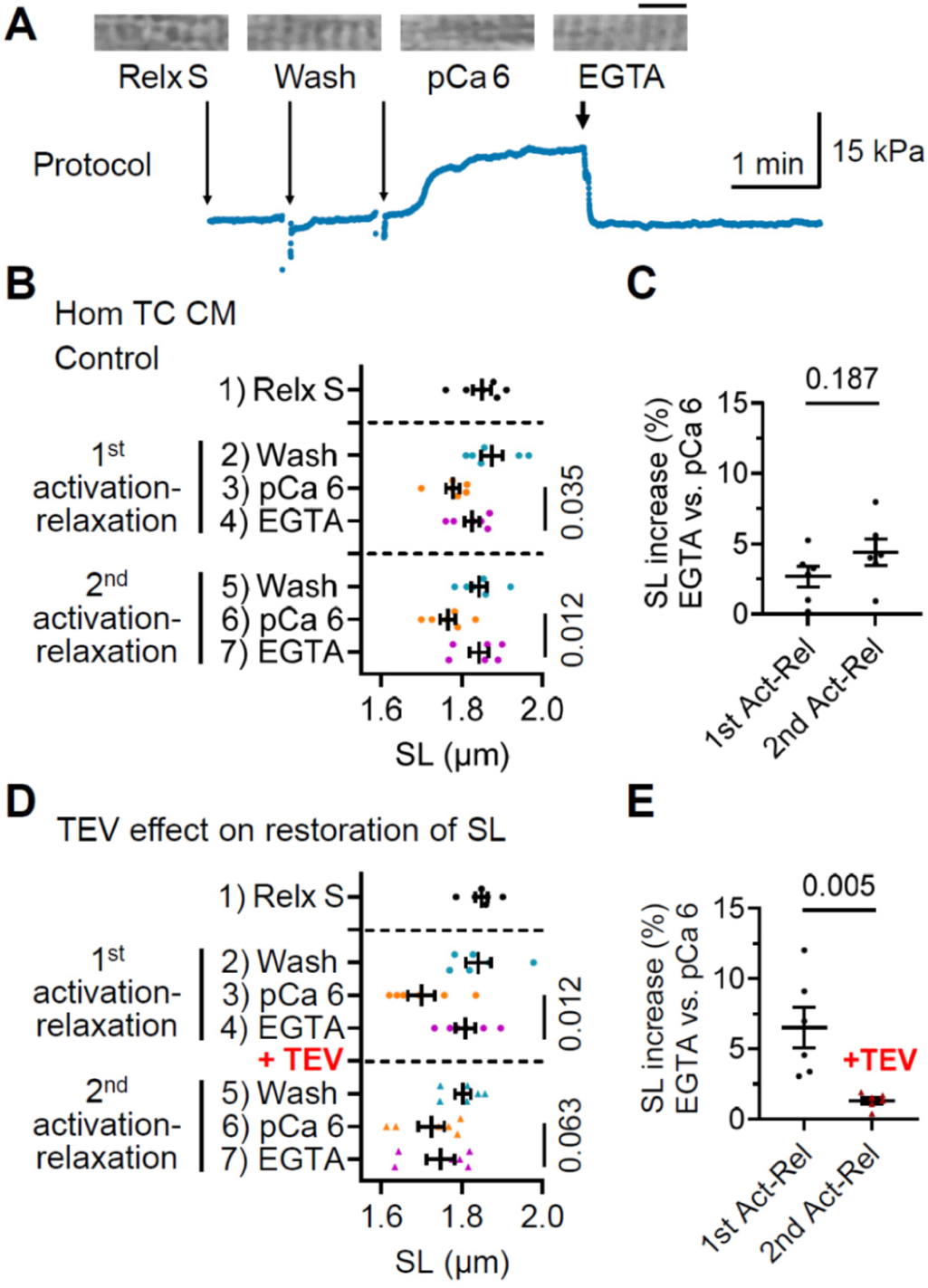
Compromised restoration of sarcomere length (SL) after contraction in titin-cleaved cardiomyocytes. **(A)** Representative force trace and phase images of sarcomeres in a single, permeabilized, Hom TC LV cardiomyocyte during an activation-relaxation protocol (activation at pCa 6; relaxation with EGTA). Scale bar, 5 µm. **(B)** Control experiments showing reproducible restoration of resting SL in successive activation-relaxation cycles in the absence of titin cleavage. Statistical significance was assessed by repeated measures one-way ANOVA with Tukey’s multiple comparisons test. **(C)** Average SL increase during EGTA-induced relaxation relative to SL during pCa 6 activation in two successive cycles of control measurements. Statistical significance was determined by an unpaired two-tailed t-test. **(D)** Absence of SL restoration with EGTA-induced relaxation following contraction in ex vivo titin-cleaved cardiomyocytes (induced by 20-minute-treatment with TEV protease), compared to pre-cleavage measurements. *Top*: repeated measures one-way ANOVA with Tukey’s test; *bottom*: Friedman test with Dunn’s multiple comparisons test. **(E)** Effect of TEV-induced titin cleavage on average SL increase during EGTA-induced relaxation relative to SL during pCa 6 activation. Statistical significance was determined by an unpaired, two-tailed t-test. All data are presented as mean ± SEM; n=6 cells/group. Relx S, relaxing solution; pCa 6, activating solution (free Ca²⁺ concentration of 10⁻⁶ mol/l); EGTA, ethylene glycol-bis(β-aminoethyl ether)-N,N,N′,N′-tetraacetic acid.

### Molecular profiling reveals altered cell-matrix crosstalk and onset of rescue mechanisms

To elucidate intramyocardial molecular changes following titin cleavage, we performed proteomic and transcriptomic analyses on LV tissues from TEV-versus GFP-injected TC mice at D6 and D13. Gene enrichment analysis with Gene Ontology (GO) and Kyoto Encyclopedia of Genes and Genomes (KEGG) revealed that significantly altered terms were consistently detected at both D6 and D13 in the proteome, whereas fewer common terms appeared at both time points in the transcriptome (**Fig. 4A; Fig. S6, Fig. S7**). This observation indicates a more stable shift in protein-level biological processes compared to transient gene expression changes. Principal component analysis clearly segregated TEV- and GFP-injected samples proteomically at both time points (**Fig. S6B-C**), with transcriptomic differences emerging only by D13 (**Fig. S7B-C**). Key proteomic GO terms at D6 included ‘collagen fibril organization’, ‘extracellular matrix organization’, ‘sarcomere organization’, ‘actin cytoskeletal organization’, and ‘cell-matrix adhesion’ (**Fig. 4A**). Volcano plots showed upregulation of collagen-organizing proteins (e.g., Col1a1, Col3a1, Loxl2, Loxl3) and select sarcomeric/cytoskeletal proteins (e.g., Myh7, Xirp 1/2, titin), with concurrent downregulation of others (e.g., Lmod3, Myo1f, Myo5b); expression of cell-adhesion molecules was also variable (e.g., increased Postn, Col8a1, Vcam1, and Itgb2; decreased Lamc2, Fbln5, and Itgb3) (**Fig. 4B-D**). Although these pathways remained enriched at both time points, additional pathways such as PI3K/PKB, TNF signaling, and mitochondrial organization emerged by D13, suggesting a compensatory response (**Fig. 4A**).

**Figure 4.**
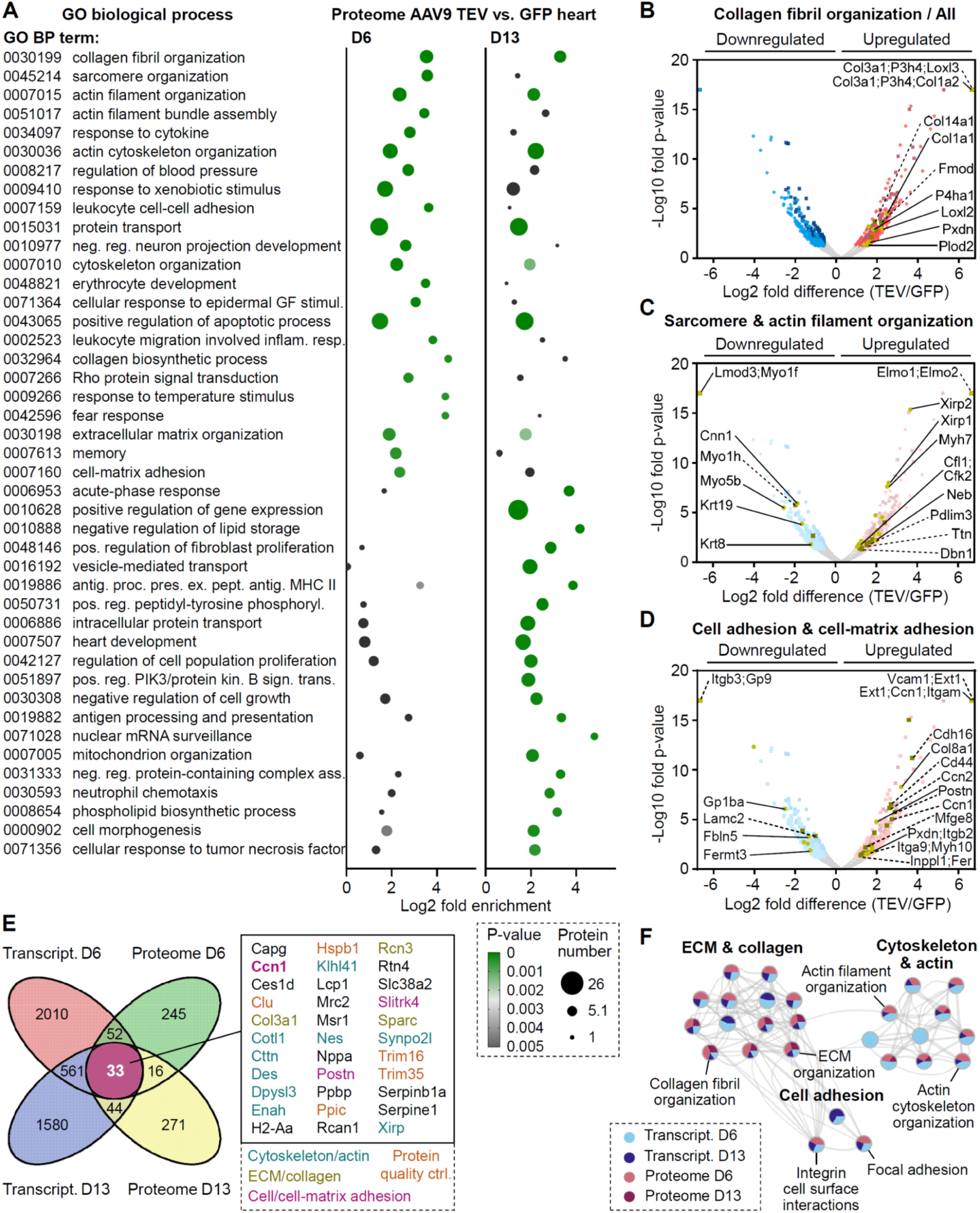
Proteomics and transcriptomics of titin-cleaved hearts. **(A)** GO biological process analysis of significantly up- or down-regulated proteins in TEV versus GFP samples, performed using DAVID. Color indicates *p*-value (threshold *p*=0.005), and circle size reflects the number of proteins. GO terms at D6 are displayed in order of increasing *p*-value and are limited to highly significant terms (*p*≤0.001); terms with *p*>0.001 are shown in dark grey. **(B-D)** Volcano plots for cardiac proteins in TEV versus GFP samples at D6 (circles) and D13 (squares) for the GO terms: (B) Collagen fibril organization (GO:0030199); (C) Sarcomere and actin filament organization (GO:0045214; GO:0007015); and (D) Cell adhesion and cell–matrix adhesion (GO:0007160; (GO:0007155). All proteins meeting the criteria of fold-change >1.5 (p<0.05) are color-coded— blue for downregulation and purple for upregulation (shown as faint color in (C) and (D)—whereas proteins designated by the respective GO terms are highlighted in yellow; select proteins are identified via solid (D6) and dashed lines (D13). **(E)** Venn diagram illustrating the overlap of significantly altered proteins/genes from the proteomic and transcriptomic datasets. The 33 common molecules are listed, with key molecules/pathways highlighted. **(F)** Network analysis combining D6 and D13 proteomic and transcriptomic data. Each circle node is a pie chart representing a GO term; the pie sections are proportional to the normalized gene count from each ‘omic’ dataset. Nodes with a similarity score >0.3 are connected. Network visualization was performed using Cytoscape. D6: N=6 (Hom TEV and Het GFP); N=3 (Hom GFP); D13: N=6 (Hom TEV and Hom GFP).

At the transcript level, enriched pathways included ‘ECM-receptor interaction’, ‘focal adhesion’, and ‘regulation of actin cytoskeleton’ at both D6 and D13 (**Fig. S7A**). Venn diagram analysis identified a core set of 33 genes and proteins commonly dysregulated at both time points, predominantly linked to cytoskeletal, ECM/collagen, and cell adhesion functions (**Fig. 4E**) – a finding reinforced by network analysis (**Fig. 4F**). Several of these molecules also fell under the ‘protein quality control’ umbrella, with ‘protein ubiquitination’ notably affected at the proteome level and several E3 ubiquitin ligases being upregulated at both time points (**Fig. S6D**). This implied activation of rescue mechanisms that also included the GO term ‘positive activation of apoptotic process’ at both D6 and D13 (**Fig. 4A**).

To independently validate these rescue pathways, we assessed apoptosis via the TUNEL assay (**Fig. S8A**), which revealed increased apoptotic activity in titin-cleaved hearts reaching statistical significance by D13 (p<0.045; **Fig. S8B**). Anti-pan-ubiquitin WBs, leveraging titin’s large size to minimize signal interference (**Fig. S8C**), indicated that cleaved (but not intact) titin was more heavily ubiquitinated in both Hom and Het LV tissues, particularly at D6 (**Fig. S8D**), suggesting rapid targeting for proteasomal degradation. Additionally, muscle E3 ligases (Atrogin-1, CHIP [Stub1], and MuRF1) were elevated in titin-cleaved Hom LV (**Fig. S8E**). In contrast, known I-band titin-binding chaperones (*31, 32*) were unchanged (HSP90, αB-Crystallin) or downregulated (HSP27) (**Fig. S8F**), while the autophagy adapter p62 was upregulated (**Fig. S8G**). Notably, these alterations were absent in TEV-versus GFP-injected Het LV tissues (**Fig. S8H-J**). Furthermore, LC3BII levels and LC3B puncta increased in D13 Hom (but not Het) TC hearts, although the LC3BII:I ratio remained unchanged, indicating limited autophagic flux (**Fig. S8K-N**). Collectively, these findings confirm that titin cleavage activates apoptosis and ubiquitin-dependent degradation as adaptive rescue mechanisms.

### Titin cleavage leads to impaired mechanical connectivity and fibroblast proliferation

Given that our omics data indicated altered cell-matrix interactions and ECM re-organization already at D6, we additionally explored these aspects via microscopic tissue analysis to elucidate the mechanistic basis of cardiac remodeling following titin cleavage. Wheat germ agglutinin (WGA) staining of tissue cross-sections (**Fig. 5A**) allowed quantitation of cardiomyocyte number and size, revealing no evidence of hyperplasia or hypertrophy (**Fig. 5B-C**). However, Ki67 staining indicated increased mitotic activity, with Ki67-positive nuclei peaking at 6% on D6 and declining to 2% by D13 (**Fig. 5A**, **D**), whereas GFP controls remained below 1%. Co-staining with Pericentriolar Material 1 (PCM1; (*33*)) and DAPI (nuclei) distinguished resting cardiomyocytes (perinuclear PCM1) from resting (PCM1-negative) and cycling (intranuclear PCM1) interstitial cells (**Fig. 5E**). Co-labeling with the activated fibroblast marker Periostin confirmed that the proliferating cells were myofibroblasts confined to interstitial niches (**Fig. 5F**). Notably, Ki67-positive cells began to rise as early as D4 when titin cleavage was still only ∼12% (**Fig. 1G**), suggesting immediate fibroblast activation in response to titin stiffness loss, preceding any overt functional phenotype.

**Figure 5.**
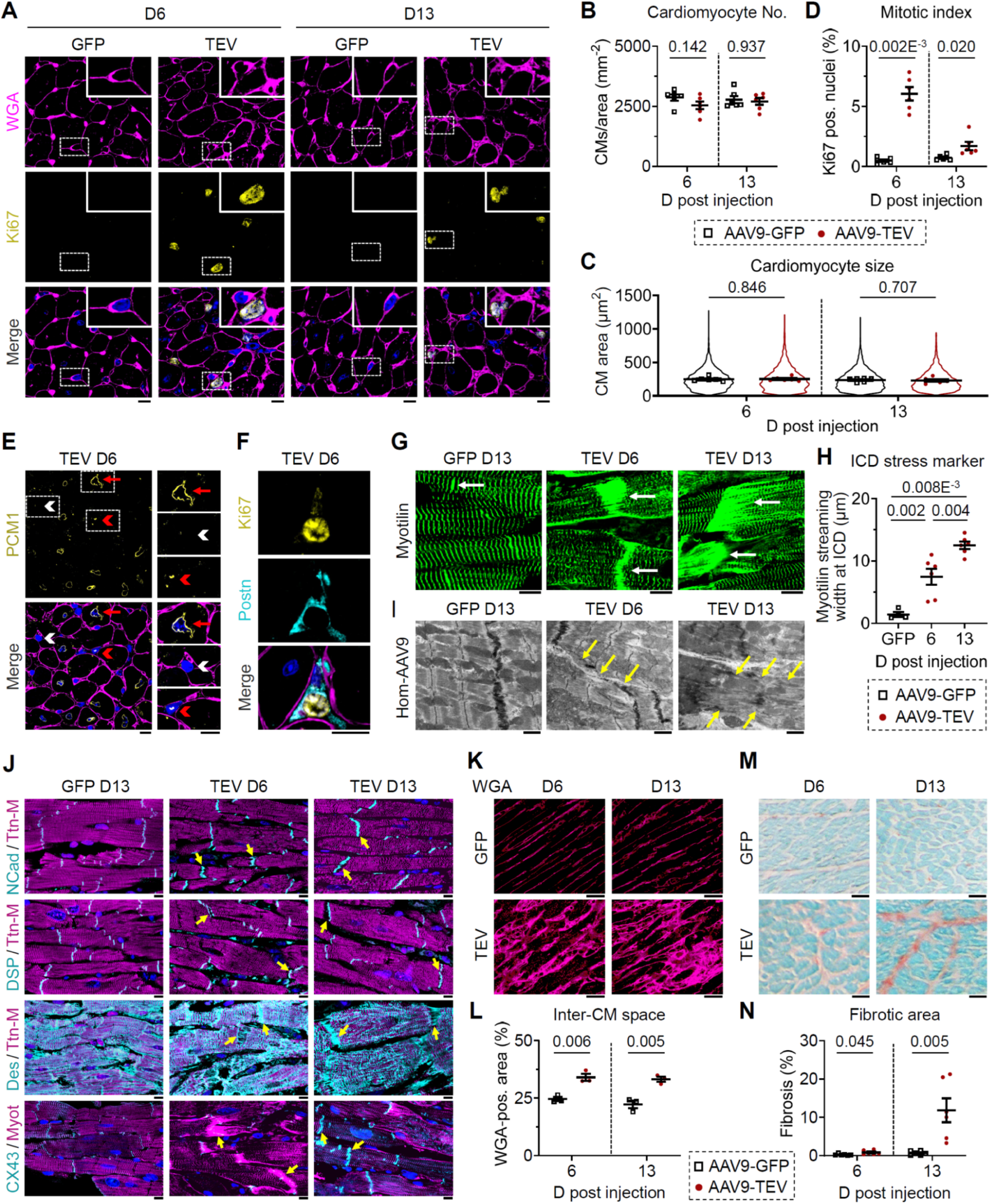
Myocardial remodeling in titin-cleaved hearts. **(A)** IF staining of Hom TC LV tissue sections at D6 and D13 post-AAV9 injection. *Top*: WGA (cell borders; magenta); *middle*: Ki67 (cycling cells; AlexaFluor 488); *bottom*: merged with DAPI (blue). Scale bars, 10 µm; insets show magnified ROIs. **(B–C)** Semi-automated quantitation of (B) cardiomyocyte number and (C) size using Nikon NIS software. Between 3,324 and 3,662 cardiomyocytes per group (N=6 mice/group); (B) violin plots show individual data distribution. **(D)** Cardiac cell proliferation (mitotic index) assessed as the percentage of Ki67-positive nuclei among 12,155-17,063 nuclei per group (N=6 mice/group). **(E)** IF staining for PCM1 (AlexaFluor 488), WGA (magenta), and DAPI (blue) at D6 post-TEV injection. Red arrows indicate resting CMs; red arrowheads, actively cycling interstitial cells; white arrowheads, resting interstitial cells. Scale bars, 10 µm. **(F)** IF images of Ki67-positive activated fibroblasts (AlexaFluor 488; *top*) co-stained for Periostin (Postn; Cy3; *middle*) and merged (*bottom*) with DAPI (blue) and WGA (magenta) at D6 post-AAV9-TEV injection. Scale bars, 10 µm. **(G)** IF images of intercalated disk (ICD) regions (arrows) in longitudinal heart sections stained with anti-Myotilin (AlexaFluor 488) from AAV9-GFP and AAV9-TEV mice at D6 and D13. Scale bars, 10 µm. **(H)** Quantitation of Myotilin signal width at ICDs (N=4 for GFP, N=6 for TEV at D6 and D13). **(I)** Transmission electron micrographs of ICDs; yellow arrows indicate ICD distortion. Scale bars, 1 µm. **(J)** IF staining for N-Cadherin (NCad; AlexaFluor 488), Desmoplakin (DSP; AlexaFluor 488), Desmin (DES; AlexaFluor 488), all co-stained with M-band titin (Ttn-M; Cy3), and Connexin 43 (CX43; AlexaFluor 488), co-stained with Myotilin (Myot; Cy3); yellow arrows highlight ICD regions. Scale bars, 10 µm. **(K)** IF images of WGA-stained Hom TC heart sections at D6 and D13 post-AAV9 injection (GFP versus TEV). Scale bars, 10 µm. **(L)** Quantitation of intercellular (WGA-positive) area as a proportion of total area in GFP- and TEV-injected groups at D6 and D16 (N=3 mice/group). **(M)** Picrosirius-red staining of Hom TC heart tissue at D6 and D13 post-AAV9 injection, comparing TEV and GFP hearts. Scale bars, 10 µm. **(N)** Quantitation of interstitial fibrosis as Picrosirius red-positive area normalized to total area (N=6 mice/group). All data are expressed as mean ± SEM; statistical significance was determined by unpaired two-tailed t-test or one-way ANOVA with Tukey’s multiple comparisons test (panel H).

We speculated that increased internal mechanical stress triggered by titin stiffness loss might drive fibroblast activation. Anti-cTEV staining showed that titin-cleaved sarcomeres lost contact with ICDs (**Fig. 1I**, arrows), while Myotilin staining demonstrated ICD-associated “streaming” (**Fig. 1I**; **Fig. 5G**). Quantitation of streaming in longitudinal LV sections showed a widened Myotilin-positive area at D6, which expanded further by D13 (**Fig. 5H**), indicative of enhanced mechanical stress. Electron microscopy confirmed progressive ICD deformations between D6 and D13 (**Fig. 5I**). Although key ICD components, such as N-cadherin (adherens junctions), Desmoplakin (desmosomes), and Connexin 43 (gap junctions), remained largely aligned, heightened signal jaggedness and disconnection of Desmin intermediate filaments implied elevated shear stress (**Fig. 5J**). Strikingly, WGA staining revealed a markedly enlarged inter-cardiomyocyte space at D6 and D13 (**Fig. 5K-L**), suggesting loss of cell-cell contact that likely contributes to LV wall thickening and volume reduction observed in titin-cleaved hearts. Finally, as anticipated from the fibroblast activation dynamics, Picrosirius-red staining demonstrated a mild rise in interstitial fibrosis at D6 and a pronounced increase by D13 (**Fig. 5M-N; Fig. S8O**), consistent with our multi-omics data. Taken together, the loss of titin stiffness and subsequent relaxation deficits compromise sarcomere-ICD connectivity and promote lateral cardiomyocyte detachment, thereby destabilizing the mechanical homeostasis sensed by fibroblasts and triggering matrix remodeling.

## Discussion

Our study provides unprecedented insights into the dual role of titin-based sarcomere stiffness in cardiac mechanics: maintaining elastic recoil and mediating cell-matrix crosstalk. By selectively cleaving elastic titin in vivo, we show that the loss of titin stiffness predominantly impairs its ability to generate restoring forces. Consequently, the heart fails to return to its diastolic shape, rapidly leading to concentric remodeling, impaired ventricular filling, and low-output heart failure. Although titin has long been recognized as a molecular spring limiting cardiomyocyte extensibility and myocardial distensibility (*5, 6*), our findings demonstrate that its role in producing restoring forces during the cardiac cycle, predicted by isolated cardiomyocyte studies (*28, 29*) but previously elusive in vivo, is even more functionally relevant. These forces drive cardiomyocyte relaxation after contraction (*34–36*) and are often compromised in disease (*37–39*). In our TC model, selective proteolysis of elastic titin traps cardiomyocytes in a contracted state, markedly reducing LV cavity size in the presence of relatively preserved systolic function. This restrictive filling pattern is accompanied by modest wall thickening, not due to hypertrophy or hyperplasia, but reflecting a loss of mechanical homeostasis as evidenced by disrupted ICD connectivity and reduced cardiomyocyte contacts, which rapidly expand the interstitial space.

A second major finding is that the cleavage of titin springs immediately activates fibroblasts, initiating a cascade of ECM remodeling events. Our multi-omics, immunofluorescence, and ultrastructural analyses indicate that the loss of titin-based stiffness and the resulting mechanical dysregulation – including increased shear stress and altered intra- and intercellular connectivity – serve as potent signals for fibroblast proliferation. These activated fibroblasts accumulate in interstitial niches and secrete collagen, triggering progressive fibrosis. While fibroblast activation in response to altered mechanical stimuli is well-established in the heart and other organs (*20, 25*), our results clearly demonstrate it as a consequence of reduced titin stiffness. Whether fibroblasts directly detect titin stiffness loss, the resulting mechanical destabilization, or both is yet to be determined.

Molecular profiling further supports that titin stiffness loss disrupts mechanical homeostasis, stimulating early compensatory pathways next to ECM remodeling, including apoptosis and protein quality control mechanisms, that appear to partially and transiently restore heart geometry and function. However, these mechanisms may eventually fail, ultimately leading to dilatation and systolic failure. Intriguingly, our experiments indicate that the heart can maintain its structural and functional integrity up to a critical threshold of ∼30% titin cleavage. This observation holds promise for therapeutic strategies targeting titin expression in conditions associated with titin proteolysis, such as acquired cardiomyopathies (*15, 17*) and atrial fibrillation (*14*), but also inherited cardiomyopathy due to I-band titin truncation (*40, 41*), suggesting that even partial preservation or restoration of titin integrity may be sufficient to maintain functional myocardial performance.

In summary, selective in vivo titin cleavage abolishes the restoring forces for elastic recoil, precipitating concentric remodeling and diastolic heart failure. Reduced titin stiffness disrupts mechanical coupling and elevates internal stress to drive fibroblast proliferation and fibrotic remodeling, demonstrating titin’s dual role in cardiac mechanical homeostasis and heart failure pathogenesis.

## Supporting information

This Excel files shows an additional summary of our proteomic analysis of TC mouse hearts injected with TEV or GFP, on D6 and D13 post injection.

The zipped file contains five Excel tables with the raw data and results of statistical tests for each main figure.

The zipped file contains eight Excel tables with the raw data and results of statistical tests for each supplementary figure.

## Acknowledgments

The authors would like to thank Lena Wildschütz for assistance with microscopy preparation, David Ing for developing the fracture-score algorithm, Nina Nagelmann for conducting MRI recordings, and Seong-won Han for help with overseeing the animals. Figure 1B was created with BioRender.com.

## Funding

We would like to acknowledge funding from

German Research Foundation grant LI 690/14-3 (WAL)

German Research Foundation grant LO2951/2-1 (CML)

Interdisziplinäres Zentrum für Klinische Forschung Münster grant Li1/012/24 (WAL)

Interdisziplinäres Zentrum für Klinische Forschung Münster support of core unit PIX (CF)

German Centre for Cardiovascular Research (DZHK) grant 81X2300190 (WAL, OJM)

Medizinerkolleg - Dean of the Medical Faculty Münster stipend 2024-25 (PH)

National Institutes of Health grant HL136737 (JAK)

National Institutes of Health grant HL172492 (JAK)

National Institutes of Health grant HL175964 (JAK)

## Author contributions

Conceptualization: WAL

Methodology: JKF, PH, CML, AU, FK, LW, SH, MMD, RH, CF, JAK, VH, OJM, WAL

Investigation: JKF, PH, CML, AU, FK, AJK, LW, SH, MMD, RH, JAK, VH

Visualization: JFK, PH, CML, AU, FK, VH, WAL

Funding acquisition: CML, CF, JAK, VH, OJM, WAL

Project administration: AH, CF, JAK, OJM, WAL

Supervision: AH, CF, JAK, OJM, WAL

Writing – original draft: WAL

Writing – review & editing: JKF, PH, CML, FK, CF, JAK, VH, OJM, WAL

## Competing interests

JAK and WAL consult and provide contract research services for multiple biotech companies, but none of these are directly related to the work performed in this study. All other authors declare no competing interests.

## Data and materials availability

All data are available in the main text or the supplementary materials. The raw mass spectrometry data has been uploaded to the MassIVE data repository (https://massive.ucsd.edu/ProteoSAFe/static/massive.jsp), a member of the ProteomeXchange Consortium (https://www.proteomexchange.org/). The data can be accessed by reviewers via private link (ftp://MSV000097567@massive-ftp.ucsd.edu, password: d3E3H0e50ghkA55z), and publicly after publication (ftp://massive-ftp.ucsd.edu/v09/MSV000097567/).

## Supplementary Materials

## Materials and Methods

### Animal model and muscle preparation

All experimental procedures were approved by the Animal Care and Use Committees of North Rhine-Westphalia, Germany (LANUV NRW, 81-02.04.2019.A472). Throughout all procedures, mice were singly housed in individually ventilated cages, maintained on a 12 hour light/12 hour dark cycle, and provided food and water ad libitum. Prior to experimentation, animals were acclimated in the local university’s animal facility, where cage occupancy varied within – but never exceeded – the maximum allowable density.

We utilized the titin cleavage (TC) mouse model, which harbors a tobacco etch virus (TEV) protease recognition site and a HaloTag cassette inserted into the elastic region of titin, near the I-band/A-band junction (*27*). These mice are healthy and exhibit a normal lifespan under baseline conditions, unless the TEV protease cleavage site is enzymatically activated (*27*). Genotyping to distinguish between homozygous (Hom), heterozygous (Het), and matched wildtype (Wt) animals was performed by PCR, as described (*9*).

To express TEV protease in the hearts of TC mice, a delivery system based on adeno-associated virus serotype 9 (AAV9) was employed. The optimized open reading frame of the TEV protease was subcloned from the plasmid pMHT238Delta into a self-complementary AAV (scAAV) genome plasmid and placed under the control of the human cardiac troponin T promoter (*Tnnt2*) to ensure cardiac-specific expression. For AAV production, the scAAV genome plasmid was co-transfected with the helper plasmid pDP9rs into low-passage HEK293T cells. AAV vectors were subsequently purified and titrated as previously described (*42*). An analogous AAV9 vector expressing enhanced green fluorescent protein (GFP) was used as a control.

Mice (both Hom and Het) received intravenous injections of 1×10^¹²^ viral particles under anesthesia (1.5-2.5% v/v isoflurane in O₂). To promote cardiac activity and induce physiological stress, mice were provided with access to a running wheel from two days prior to injection until 13 days post injection; during this time, they were hosted individually. Transthoracic echocardiography (TTE) was performed on the day of injection (D0) and subsequently twice per week. Cardiac magnetic resonance imaging (cMRI) was conducted at day (D)6 and D13 post injection. Mice aged 9-44 weeks were assigned to TTE, whereas those selected for cMRI ranged from 8-18 weeks of age. For echocardiography, cohort sizes included 15 Hom TEV, 15 Het TEV, 15 Hom GFP, and 15 Het GFP animals. MRI group sizes were limited to 12 Hom TEV, 12 Hom GFP, 6 Het TEV, and 6 Het GFP animals. Both male and female mice were included. At either D6 or D13 post injection, and in some cases at D4, D5, or D10, mice were sacrificed by cervical dislocation. Hearts were excised, washed in cold PBS, and either flash-frozen in liquid nitrogen for biochemical assays or retrogradely perfused for further processing for electron microscopy and confocal immunofluorescence microscopy.

### Transthoracic echocardiography

For ultrasound examinations, mice were anesthetized with 1.5-2.5% v/v isoflurane/O2, and the chest area was shaved with depilatory gel. Imaging was performed according to established protocols (*43*) using a Vevo 2100 (VisualSonics) ultrasound system (18–55 MHz transducer, MS550D). Heart rate (under anesthesia) was continuously recorded during the experiment. Left ventricular (LV) structure and function were assessed via parasternal short-axis (SAX) M-mode imaging. LV posterior wall (LWPW) and interventricular septum (IVS) were directly measured in both systole and diastole. LV end-systolic volume (LVESV) and LV end-diastolic volume (LVEDV) were derived from LV Trace. From these, LV ejection fraction (LVEF), fractional shortening (FS), stroke volume (SV), and cardiac output (CO) were calculated as key indicators of ventricular performance. Pulsed-wave (PW) Doppler imaging of the mitral valve was used to measure Emax and Amax, from which the E/A ratio was determined. Pressure half-time (PHT) and acceleration (Ac) were also assessed. Tissue Doppler imaging (TDI) of the medial and lateral mitral annulus was performed, and e’ and a’ were extracted from the TDI curve, with corresponding ratios calculated. PW Doppler imaging of the aortic valve was used to evaluate LV ejection. All parameters were analyzed with Vevo Lab 2.2.0 software.

### Cardiac magnetic resonance imaging

For cardiac magnetic resonance imaging (cMRI), mice were placed in a warmed animal bed and 1-2% v/v isoflurane/O^2^ was administered via a breathing mask. Breathing rate and body temperature were continuously monitored during measurements, and anesthesia depth was adjusted as needed. In vivo cMRI was performed at 9.4 T using a Bruker BioSpec 94/20 system (Ettlingen, Germany) with a 1 T/m gradient system and ParaVision 5.1 software, including IntraGate for sequence acquisition and reconstruction. A 35 mm volume coil was used for data acquisition. Systolic and diastolic heart function were assessed by acquiring a stack of contiguous short-axis (SAX) slices (1 mm thickness) covering the entire right and left ventricles using the self-gated cine FLASH (IntraGate FLASH) sequence, as described (*44*). For analysis, regions of interest in MRI FLASH images were manually selected on the end-diastolic and end-systolic frames of each SAX-slice by tracing the epicardial and endocardial borders, ventricular and heart diameters, and septal and posterior wall thickness using ImageJ 1.54 software (NIH Bethesda). To obtain LVEDV and LVESV, the volume for each frame was calculated as the sum of the area of interest in each slice multiplied by the slice thickness. From these, SV, CO, and LVEF were derived. Outer diameter (OD) was determined in short-axis view and the Green-Lagrange strain (*E*) was quantified from the cyclic deformation of the myocardium at the mid-ventricular level during systole, based on the circumferential dimensions of the endocardial contour measured at end-diastole (CD) and end-systole (CS), using the following equation:

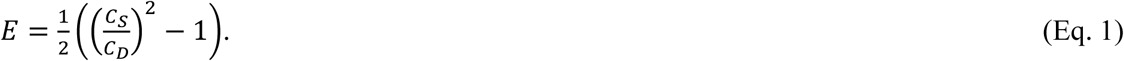

### Immunofluorescence and electron microscopy

Immunofluorescence (IF) and transmission electron microscopy (TEM) were done according to published protocols (45). For IF microscopy, LV tissue samples were fixed in 4% paraformaldehyde (PFA) and 15% saturated picric acid, followed by dehydration and paraffin embedding. Sections of 5-7 µm thickness were rehydrated, treated with peroxidase buffer and citrate-EGTA, and blocked with 5% bovine serum albumin (BSA) containing 0.5% Triton X-100 for 1 hour. The sections were then incubated with primary antibodies overnight at 4°C, followed by secondary antibody incubation under the same conditions. Detailed information on the primary and secondary antibodies is provided in Table S1.

To visualize cleaved titin at D6 and D13 following AAV9 injection, an anti-TEV-cleavage-site (cTEV) antibody was employed. Titin cleavage was quantified by normalizing cTEV signals to the corresponding anti-Myotilin signals – which serve as markers for both Z-disks and intercalated disks – using NIS Elements v. 4.3 software. The number of sarcomeres was determined by analyzing confocal images of anti-Ttn-M staining, where the M-band signals were semi-automatically counted using NIS Elements 4.3 as a measure of the sarcomere number. The width of the intercalated disks labeled with anti-Myotilin was measured using ImageJ v. 1.54. Wheat germ agglutinin (WGA) staining on tissue cross-sections delineated cell borders, facilitating the semi-automated quantification of cardiomyocyte number per unit area and cardiomyocyte size with NIS Elements 4.3 software. Cycling cells were immunostained with anti-Ki67, while the nuclei were counterstained with DAPI; the mitotic index was calculated as the percentage of Ki67-positive nuclei relative to the total number of DAPI-stained nuclei. Cardiomyocytes were identified using anti-Pericentriolar Material 1 (PCM1), whereas myofibroblasts were distinguished using anti-Periostin (Postn). The tissue area occupied by WGA staining on longitudinal sections was quantified using NIS Elements 4.3 software and expressed relative to the total tissue area. Relative LC3B expression was assessed with an anti-LC3B antibody and normalized to anti-Myotilin signals. Apoptosis was characterized using the TUNEL assay, with the apoptotic index calculated as the percentage of TUNEL-positive cells relative to the total number of DAPI-stained nuclei.

For TEM, tissue samples fixed in 4% paraformaldehyde (PFA) were longitudinally sectioned into 50-μm-thick slices using a VT 1000S Leica vibratome and washed twice in PBS. Samples were then dehydrated and embedded in Durcupan resin (Fluka). Resin blocks were prepared and ultrathin sections cut using a Leica Ultracut S microtome. The sections were mounted onto glow-discharged, Formvar-carbon-coated copper grids. Electron microscopy was performed using a Zeiss LEO 910 transmission electron microscope, and images were captured with a TRS SharpEye CCD camera (Troendle).

### Quantitation of Z-disk disorder

To quantify the level of Z-disk disorder (‘fracture score’) in AAV9-TEV-injected Het TC mice and AAV9-GFP-injected control mice at D13 post injection, we employed a semi-automatic program previously developed by us (*30*). Briefly, transmission electron micrographs of myofibers were filtered to extract Z-disk structures. Each pixel along a Z-disk line was assigned x- and y-coordinates, which were then fitted to a best-fit straight line using linear regression. The residuals were weighted based on the angular differences between regressions for all Z-disks in the x-y coordinate space, averaged, and further analyzed to determine the fracture score.

### Immunohistochemistry

Ventricular tissue samples were fixed in 4% PFA and 15% saturated picric acid, processed and embedded in paraffin. Sections of 5-7 µm thickness were rehydrated and stained with a Picrosirius-Red-Fast-Green kit to label collagen fibers. Interstitial cardiac fibrotic area was quantified as volume fraction of collagen using Image J v. 1.54.

### SDS-PAGE and immunoblotting

Standard 10-15% SDS-PAGE gels, agarose-strengthened, loose (1.8%) SDS-PAGE titin gels, and 1.8/7.5% two-phase (‘stacked’) gels, were prepared according to previously published protocols (*41*). Proteins were visualized by Coomassie staining or identified by Western blot (WB), as described (*9*, *46*). Relative band intensities on Coomassie-stained gels for intact and cleaved N2BA/N2B titin were used to quantify the percentage of total titin cleaved by TEV. Antibodies used for WB are listed in **Table S1**. Signals from HRP-conjugated secondary antibodies were visualized by chemiluminescence (Amersham ECL start WB detection reagent, GE Healthcare) and recorded using the ImageQuant LAS 4000 Imaging System (GE Healthcare). Signal intensity of detected bands was quantified by densitometry using either MultiGauge v. 3.0 (Fuji) or ImageQuant TL v. 7.1 (GE Healthcare) software.

### Mechanical measurements on isolated cardiomyocytes

Cardiomyocytes were isolated from hearts that were first washed in PBS, then flash-frozen in liquid nitrogen, and stored at -80°C until use. A small piece of ventricular tissue was excised from the frozen heart and thawed in relaxing solution (170 mM K-propionate, 20 mM MOPS, 2.5 mM Mg-acetate, 5 mM K₂EGTA, 2.5 mM ATP, 14.5 mM creatine phosphate, 1× working concentration of protease inhibitor cocktail (Promega: G6521), pH 7.0 at 4°C). The tissue was mechanically disrupted and permeabilized in relaxing solution supplemented with 0.5% Triton X-100 for 8 minutes, followed by three washes in relaxing solution. Cardiomyocytes were mechanically analyzed as described (*9*). Cardiomyocytes for mechanical measurements were obtained from altogether four Hom TEV, eight Het TEV, and four wildtype mouse hearts.

To determine the restoration of sarcomere length (SL) after contraction, 200 µL of the cardiomyocyte suspension in relaxing solution was pipetted onto a glass coverslip. Once the cells adhered to the glass by gravity, the solution was replaced with washing solution (185 mM K-propionate, 20 mM MOPS, 2.5 mM Mg-acetate, 2.5 mM ATP, pH 7.0 at room temperature), followed by activation with pCa 6 solution (170 mM K-propionate, 10 mM MOPS, 2.42 mM Mg-acetate, 31.60 mM CaEGTA, 1.72 mM K₂EGTA, pH 7.0 at room temperature). Contraction was stopped by adding 20 µL K₂EGTA (50 mM). The solution was then changed to relaxing solution, and in-house produced TEV protease (*9*) was added to cleave titin. For control measurements, cells were incubated in relaxing solution without TEV protease. After 20-30 minutes at room temperature, the solution was replaced with washing solution, and a second contraction was started by activation with pCa 6 solution, followed by EGTA-induced relaxation. SL was measured throughout using 901D Hi-Speed Video Sarcomere Length v. 4.195 (Aurora Scientific) software.

To measure the passive elastic force of single Het TC cardiomyocytes, permeabilized cells were mounted between a piezoelectric motor and a force transducer (Aurora Scientific, 403A) using shellac dissolved in 70% ethanol (120 mg/mL) on the stage of an Axiovert 135 inverted microscope (Carl Zeiss). Cells were stretched from slack length (∼1.8 µm SL) to ∼2.1 µm SL, held for 7 seconds, and then returned to slack length. Elastic (quasi-steady state) force was calculated before and after TEV protease treatment by fitting the force-relaxation segment of the raw traces with a one-phase exponential decay function in GraphPad Prism v. 8 software (*9*).

To compare the active force of Wt and Het TC cardiomyocytes before and after titin cleavage, permeabilized cells were mounted between a piezoelectric motor and a force transducer (Aurora Scientific, 403A) in relaxing solution and adjusted to ∼1.9 µm SL. The solution was then sequentially replaced with washing solution and pCa 5 solution (170 mM K-propionate, 10 mM MOPS, 2.40 mM Mg-acetate, 46.20 mM CaEGTA, 0.22 mM K₂EGTA, pH 7.0 at room temperature). Contraction was stopped by adding 20 µL K₂EGTA (50 mM). Active force was determined as the difference between peak force in pCa 5 solution and the force after EGTA addition. Cell diameter was measured using 901D Hi-Speed Video Sarcomere Length v. 4.195 (Aurora Scientific) software, and force per area was calculated, assuming a circular cross-sectional area. The active force per area of Het TC cardiomyocytes after titin cleavage was normalized to that of Wt cardiomyocytes after TEV protease incubation.

### Quantitative proteome analysis

Proteomic analyses were performed using the following number of animals: at D6 post injection, N=6 (Hom TEV). N=6 (Het GFP), N=3 (Hom GFP); and at D13, N=6 (Hom TEV and Hom GFP). For the analysis at D6, HET GFP and Hom GFP were pooled, as they showed only few differences.

#### In-solution digestion and peptide purification

Whole heart tissue was mechanically homogenized in 9 M urea containing 0.3% Triton X-100 and protease/phosphatase inhibitor cocktails (Halt). Samples were sonicated and centrifuged at 17,000 × g for 5 minutes, and the pellet of insoluble proteins was discarded. To remove Triton, 100 μg of lysate was subjected to chloroform–methanol precipitation and subsequently resuspended in 100 μL of 9 M HPLC-grade urea. Samples were reduced with 5 mM dithiothreitol (DTT) for 45 minutes at room temperature, followed by alkylation with 10 mM iodoacetamide (IAA) for 30 minutes in the dark at room temperature. Samples were then diluted with 450 μL of 50 mM HPLC-grade Tris-HCl to reduce the urea concentration to <2 M. Trypsin/Lys-C Protease Mix (2 µg; Thermo Fisher, #A41007) was added, and samples were incubated at 37°C with shaking at 600 RPM for 16-18 hours. Digestion was terminated by acidification with trifluoroacetic acid (TFA) to a final pH ≤ 3. Peptides were then dried using a vacuum centrifuge and desalted with C18 Spin Columns (G Biosciences) following the manufacturer’s instructions.

#### Liquid chromatography (LC)

Desalted peptides were again dried via vacuum centrifugation and resuspended in Buffer A (0.1% formic acid in HPLC-grade water). Peptides were loaded onto a Vanquish Neo UHPLC system (Thermo Fisher) operating in a heated trap-and-elute workflow using a C18 PepMap trap column (5 mm; P/N 160454) in forward-flush configuration. This was coupled to a 50 cm Easy-Spray™ PepMap™ Neo 2 μm C18, 75 μm×500 mm analytical column (P/N ES75500PN), with the column oven maintained at 40°C and 100% Buffer A. Peptide separation was achieved using a 150-minute gradient with Buffer B (80% acetonitrile, 0.1% formic acid): 4-5% over 5 minutes, 5-35% over 110 minutes, 35-65% over 25 minutes, 65-99% over 3 minutes, and held at 99% for 12 minutes. Mass spectra were acquired on an Orbitrap Eclipse Tribrid mass spectrometer with a FAIMS Pro interface (Thermo Fisher), running Tune 3.5 and Xcalibur 4.5 software. The spray voltage was set to 1600 V, ion transfer tube temperature to 300°C, and FAIMS compensation voltages (CVs) were cycled between -45 V, -55 V, and -65 V with a 1.5 s cycle time.

MS1 spectra were acquired in the Orbitrap at 120,000 resolution with a scan range of 375-1600 m/z, normalized AGC target of 300%, maximum injection time of 50 ms, and S-lens RF level of 30. Source fragmentation was disabled, and data were acquired in positive profile mode. Monoisotopic precursor selection was enabled (peptides only), and precursor ions with charge states of 2-7 were included (unassigned charge states were excluded). Dynamic exclusion was enabled (n=1) with a 60 s exclusion duration and ±10 ppm mass tolerance. Data-dependent MS2 (DDMS2) scans were acquired using quadrupole isolation (1.6 m/z window), higher-energy collisional dissociation (HCD) at 30% collision energy, and ion trap detection at Turbo scan rate. The AGC target was set to 10,000, maximum injection time to 35 ms, with one microscan per spectrum, and data were recorded in centroid mode.

#### MS/MS data analysis and processing

Raw data were analyzed using Proteome Discoverer 2.5 (Thermo Fisher) with the Sequest HT search engine. Spectra were searched against the *Mus musculus* UniProt protein database (uniprot-proteome_UP000000589). Search parameters included a precursor mass tolerance of 10 ppm and a fragment mass tolerance of 0.02 Da. Two missed trypsin cleavages were allowed. Variable modifications included oxidation (Met) and N-terminal acetylation; carbamidomethylation (Cys) was set as a static modification. Peptide spectral match (PSM) validation was performed using Percolator with a strict false discovery rate (FDR) of 0.01, a relaxed FDR of 0.05, a maximum ΔCn of 0.05, and validation based on q-values. For precursor ion quantitation, Unique + Razor peptides were used. Precursor abundance was determined by intensity, normalized to total peptide amount. Protein abundance was calculated by summing the intensities of connected peptides, and protein ratios were derived from protein abundances.

The raw mass spectrometry data has been uploaded to the MassIVE data repository (https://massive.ucsd.edu/ProteoSAFe/static/massive.jsp), a member of the ProteomeXchange Consortium (https://www.proteomexchange.org/). The data can be accessed by reviewers via private link (ftp://MSV000097567@massive-ftp.ucsd.edu, password: d3E3H0e50ghkA55z), and publicly after publication (ftp://massive-ftp.ucsd.edu/v09/MSV000097567/).

#### Bioinformatics and data visualization

Principal component analysis (PCA) was performed using Perseus software (v. 1.6.15.0). Gene ontology (GO) and Kyoto Encyclopedia of Genes and Genomes (KEGG) pathway enrichment analyses were conducted using DAVID v2021 (knowledgebase v2023q4) (*47*, *48*), with the default *Mus musculus* gene list as background.

Data visualization was carried out using GraphPad Prism 9 (e.g., volcano plots, GO biological processes), and R (v. 4.4.2) with the ggplot2 package (for combined GO BP, MF, CC, and KEGG visualizations). Venn diagrams were generated using the online tool from the Van de Peer laboratory (http://bioinformatics.psb.ugent.be/webtools/Venn). For network analysis, refer to the transcriptome analysis section.

### Transcriptomics

RNA extraction, library construction, sequencing and bioinformatics analysis were performed by Novogene (Munich, Germany). Briefly, to prepare LV tissue for RNA sequencing, ∼10 mg tissue was collected at D6 and D13 post injection, flash-frozen and sent to Novogene, for RNA isolation and sequencing. RNA was extracted using a trizol based method. Preliminary quality control was performed on 1% agarose gels, to test for RNA degradation and potential contamination. Sample purity, RNA integrity, and final expression were measured using Bioanalyzer 2100 (Agilent Technologies, USA). For Library construction, quality control and sequencing, eukaryotic rRNA removal kit was used to remove ribosomal RNA, and rRNA-free residues were cleaned up by ethanol precipitation. Subsequently, sequencing libraries were generated using the rRNA-depleted RNA. After fragmentation, the first strand of cDNA was synthesized using random hexamer primers. Then the second strand cDNA was synthesized and dUTPs were replaced with dTTPs in the reaction buffer. The directional library was ready after end repair, A-tailing, adapter ligation, size selection, USER enzyme digestion, amplification, and purification. Quantified libraries were pooled and sequenced on the Illumina Novaseq X plus platform using V1.5 reagent, S4 flow cell with paired-end strategy PE150. Sequencing depth was 40 million reads.

#### Transcriptomics bioinformatics

Bioinformatics Analysis was conducted by Novogene, China. Briefly, quality control raw data (raw reads) in fastq format were initially processed to obtain clean data (clean reads) and all downstream analyses were performed based on the clean data with high quality. Next, the reference genome, ensembl_100_mus_musculus_grcm38_toplevel, and gene model annotation were obtained, and an index of the reference genome was built using Hisat2 v2.0.5; paired-end clean reads were aligned to the reference genome. Gene expression was then quantified according to company standards.

Differential expression analysis of TEV vs. GFP injected samples was performed using the DESeq2 R package (1.20.0). For D6 post injection samples, Hom GFP (N=3) and Het GFP (N=6) were pooled and compared with Hom TEV (N=6); for D13 post injection samples, Hom TEV (N=6) and Hom GFP (N=6) were compared. The *p*-values were adjusted using the Benjamini and Hochberg’s approach for controlling the false discovery rate. GO enrichment analysis and KEGG pathways of differentially expressed genes were implemented by the ClusterProfiler (3.8.1) R package, in which gene length bias was corrected. Corrected *p*<0.05 was considered significantly enriched by differential expressed genes. Data visualization was performed using GraphPad Prism or in R package using ggplot2.

#### Network analysis

Differentially expressed gene lists of TEV vs. GFP hearts at D6 and D13, from both transcriptomic and proteomic analysis, were put through Metascape (https://metascape.org (*49*) and the network diagrams were visualized in Cytoscape. Each term was represented by a circle node pie graph where each pie section is proportional to the gene count originating from the gene list obtained from the various ‘omics’ analyses. To overcome the disproportionate number of hits seen in transcriptomics compared to proteomics, the gene count for each node was normalized to the total genes detected for that given ‘omic’ analysis. Terms with a similarity score >0.3 were linked by a line.

### Statistical analysis

Data organization, graphing, and statistical analyses were performed using Microsoft Excel 2021 and GraphPad Prism 8 or 10 software. Results are usually presented as mean ± SEM or as percentages; occasionally also as median and 95% confidence interval (CI). Normality was assessed using, at a minimum, the Shapiro-Wilk test. Significant differences between two groups were determined using a two-tailed, paired or unpaired, t-test for normally distributed data or the Mann-Whitney test for non-normally distributed data. Significant differences between more than two groups were determined using ANOVA with post hoc Tukey’s tests, or Kruskal-Wallis with Dunn’s test, as appropriate. Differences with p<0.05 were considered statistically significant.

---

**Fig. S1.**
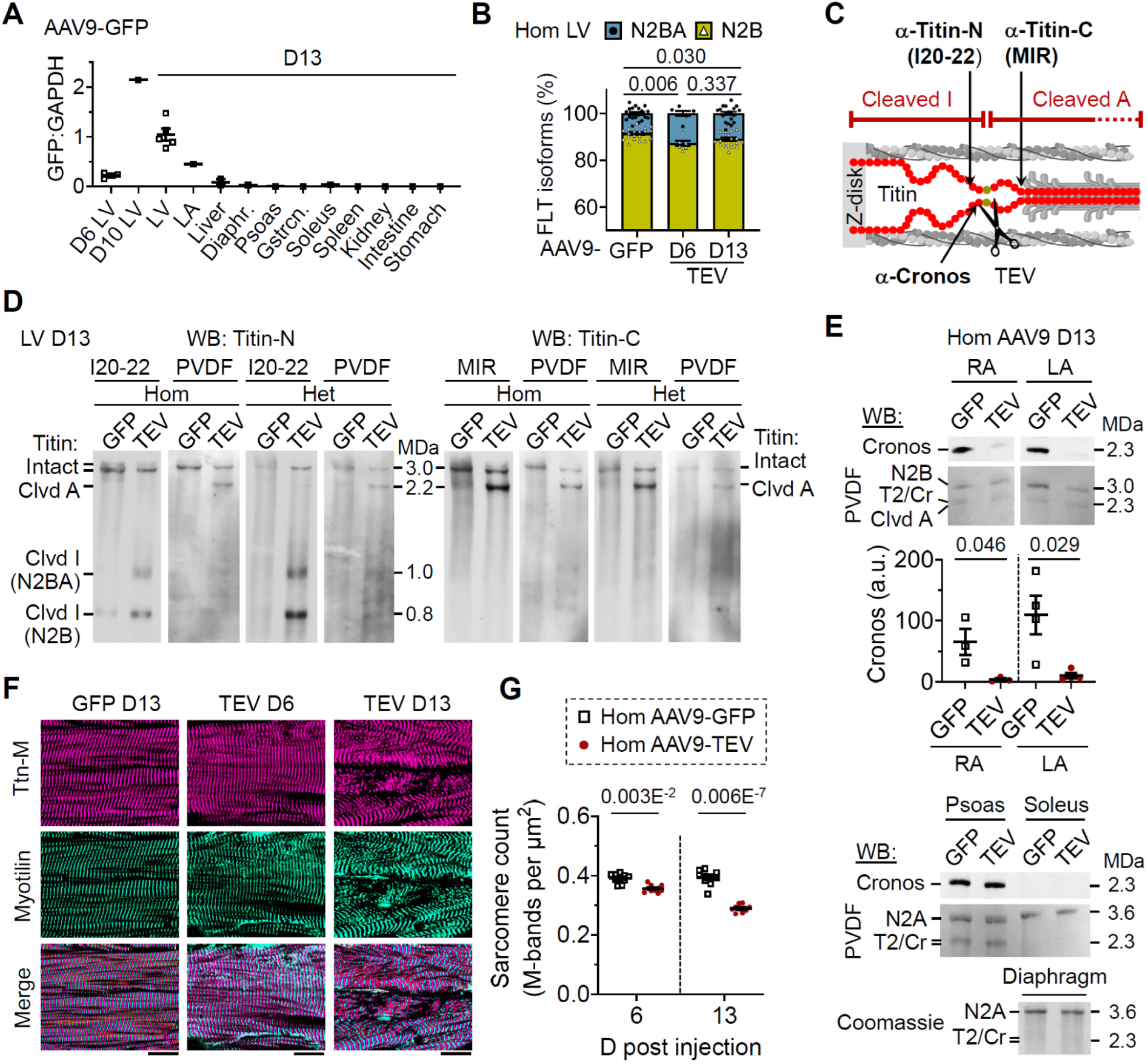
Additional cardiac protein analysis after in vivo titin cleavage. **(A)** Western blot analysis of GFP expression in various organs of AAV9-GFP-injected mice, normalized to GAPDH. Samples: D6 LV (N=3); D10 LV (N=1); D13 LV (N=5); D13 left atrium (LA), liver, diaphragm, psoas, gastrocnemius, soleus, spleen, kidney, intestine, and stomach (N=1 each). **(B)** Average titin isoform composition at D13 post-AAV9-GFP injection versus D6 and D13 post-AAV9-TEV injection. N=19 (GFP), N=6 (D6 TEV), N=19 (D13 TEV); significance by one-way ANOVA with Tukey’s multiple comparisons test. **(C)** Schematic illustrating TEV-induced titin cleavage into I-band and A-band fragments within the half-sarcomere, with the epitope locations of anti-titin antibodies (I20-22, MIR, Cronos) indicated. **(D)** Western blot confirmation of in vivo titin cleavage in TC mouse LV tissue at D13 post-TEV injection versus GFP controls, using antibodies against titin epitopes N-terminal (I20-22; *left*) and C-terminal (MIR; *right*) to the TEV-cleavage site. Analyses were performed in both Hom and Het TC hearts; PVDF staining served as a loading control. **(E)** Cronos protein expression in right atrium (RA) and LA of AAV9-GFP and AAV9-TEV Hom TC mice at D13, with PVDF as loading control. Graph: densitometric analysis (RA, N=3; LA, N=4). Bottom panels demonstrate absence of titin cleavage in skeletal muscles (WB for Cronos and Coomassie staining). **(F)** Representative confocal images of Hom TC LV tissue: *left*, D13 AAV9-GFP; *center*, D6 AAV9-TEV; *right*, D13 AAV9-TEV. *Top*: titin C-terminus (anti-TTN-M, Cy3); *middle*: Z-disks (anti-Myotilin, AlexaFluor 488); *bottom*: merge. Scale bars, 10 µm. **(G)** Semi-quantitative analysis of sarcomere counts based on TTN-M staining (NIS software); n=10 images per group; N=6 (GFP) and N=3 (TEV). All data are mean ± SEM. Except in (B), significance was determined by unpaired two-tailed t-test or Mann-Whitney test.

**Fig. S2.**
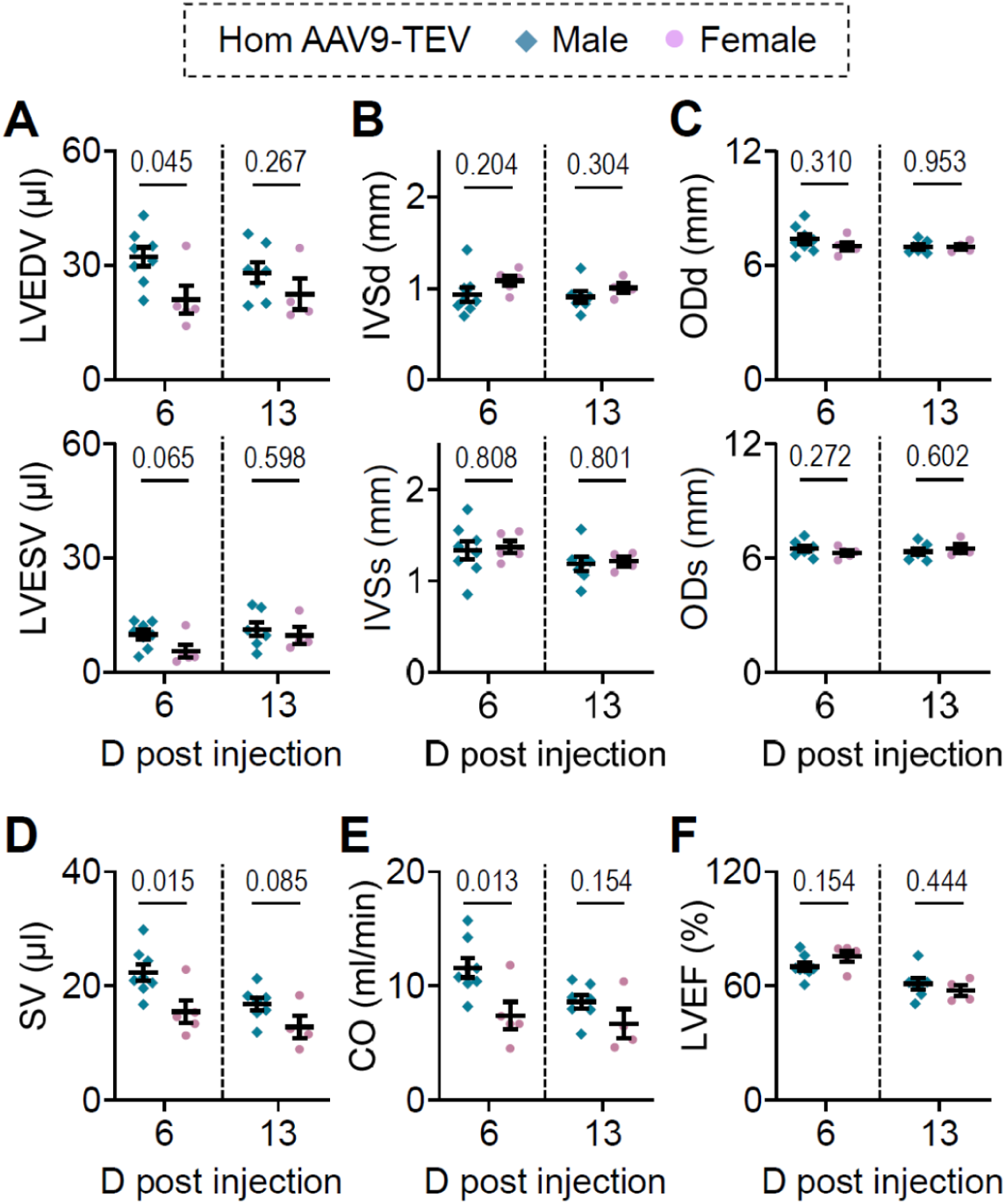
Sex-stratified cMRI analyses of AAV9-TEV-injected Hom TC mouse hearts. (**A**) LV volumes (LVEDV and LVESV); (**B**) Intraventricular septum widths (IVSd and IVSs); (**C**) Outer heart diameters (ODd and ODs); and functional parameters: (**D**) Stroke volume (SV), (**E**) Cardiac output (CO), and (**F**) LVEF. All data are mean ± SEM; N=7-8 (TEV, male) and N=4-5 (TEV, female). Statistical significance was determined by unpaired two-tailed t-test.

**Fig. S3.**
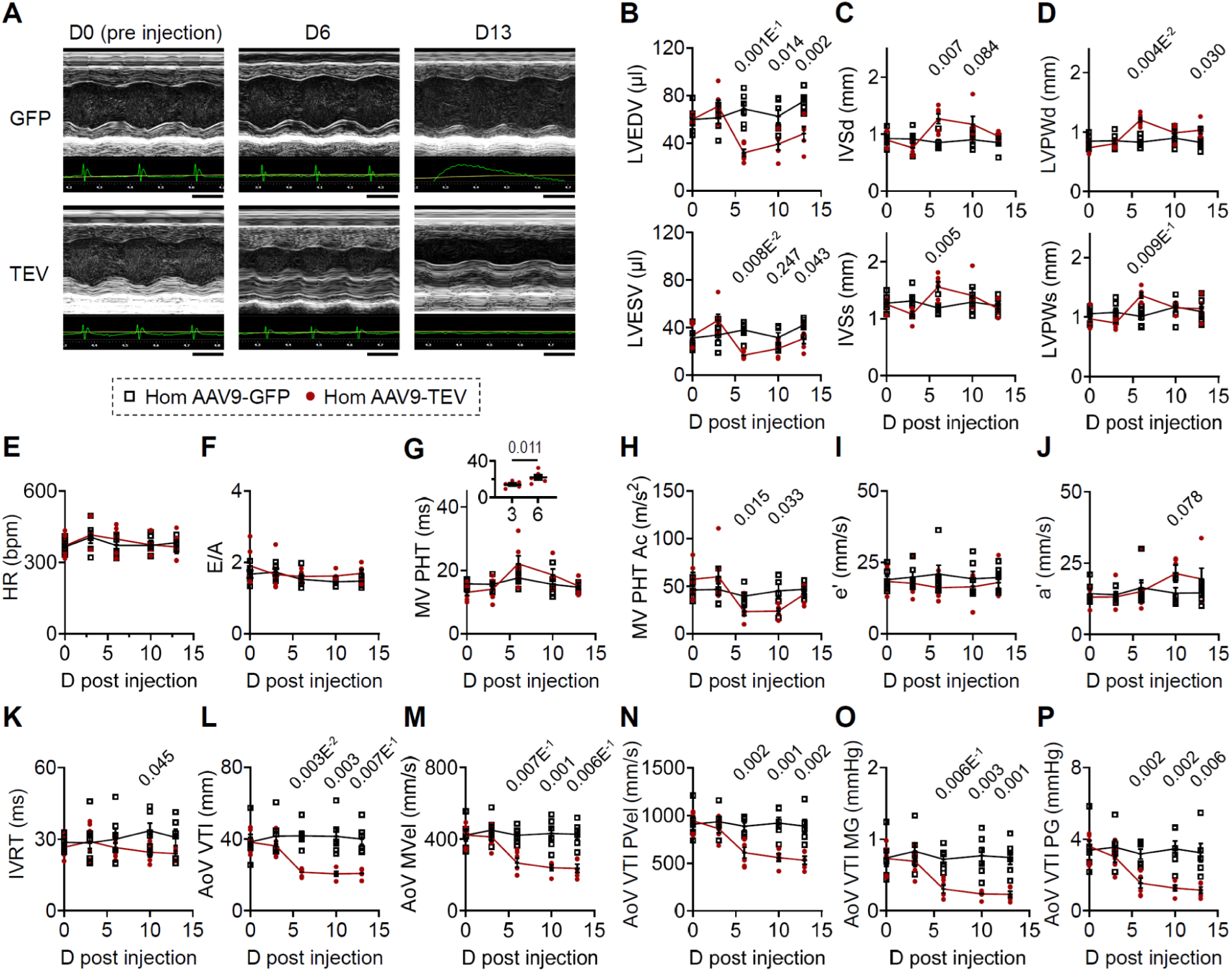
Additional transthoracic echocardiography analyses in AAV9-injected Hom TC mouse hearts. **(A)** Representative M-mode TTE images used to quantify LV volume, IVS width, and posterior wall (PW) thickness at four time points post-AAV9 injection (D0 = pre-injection). Scale bars, 0.1 s. **(B)** LV volumes: end-diastolic (LVEDV) and end-systolic (LVESV). **(C)** IVS dimensions: diastolic (IVSd) and systolic (IVSs). **(D)** LV posterior wall: diastolic (LVPWd) and systolic (LVPWs). **(E)** Heart rate (HR) under anesthesia. **(F–H)** PW-Doppler assessment of mitral valve (MV) flow: (F) E/A ratio; (G) MV pressure half time (PHT) (small graph: comparison at D3 vs. D6 in AAV9-TEV mice); (H) MV PHT acceleration (Ac). **(I–J)** Tissue Doppler imaging: (I) e′ and (J) a′ velocities. **(K)** Isovolumic relaxation time (IVRT). **(L–P)** PW-Doppler measurements of aortic valve (AoV) flow: (L) AoV velocity time integral (VTI); (M) AoV mean velocity (MVel); (N) AoV peak velocity (PVel); (O) AoV mean gradient (MG); and (P) AoV peak gradient (PG); All data are presented as mean ± SEM; N=5-6 mice/group. Statistical significance was determined by unpaired two-tailed t-test or paired t-test (small graph in (G)).

**Fig. S4.**
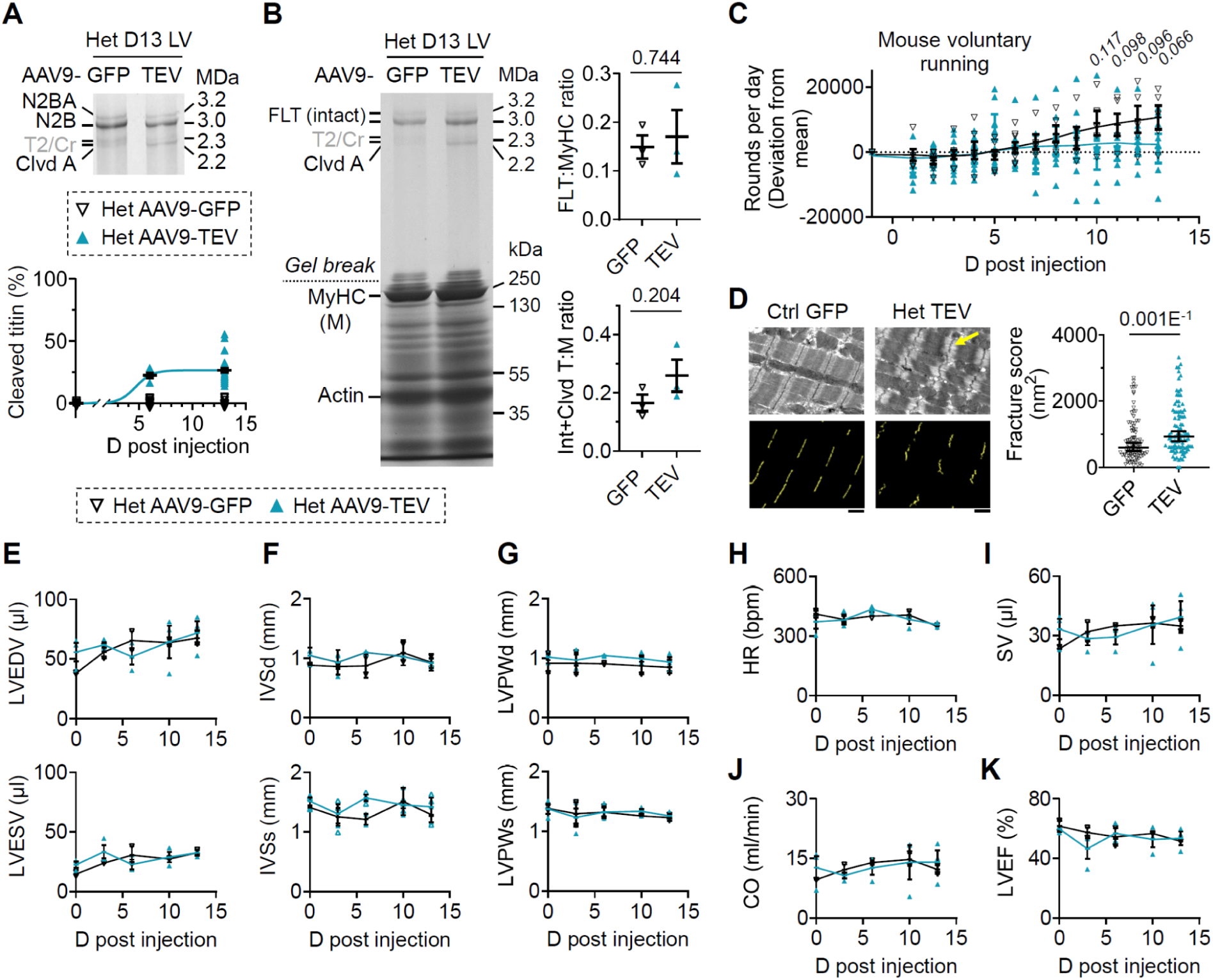
Consequences of in vivo titin cleavage in heterozygous (Het) TC mouse hearts. **(A)** Representative Coomassie-stained titin gel of Het TC mouse LV tissue at D13 post-AAV9 injection and time course of cleaved titin increase in TEV-versus GFP-injected hearts (N=9 for GFP and TEV at D13; N=6 for TEV at D6). **(B)** *Left*: Representative two-phase polyacrylamide (PA) gel (1.8% PA for titin, 7.5% PA for other cardiac proteins) from Het TC mice at D13 post-AAV9-GFP or AAV9-TEV injection. *Right*: Densitometric analysis showing the ratio of full-length (or total, intact+cleaved) titin to MyHC (M) (N=3 mice/group). **(C)** Voluntary running activity showing daily lap count (deviation from start) in Het TC mice following AAV9-GFP or AAV9-TEV injection throughout the experiment (N=6-8 mice/day for GFP; N=13-16 mice/day for TEV). **(D)** Representative transmission electron micrographs of D13 GFP- and TEV-injected Het TC hearts. Arrow indicates wavy Z-disks; scale bars, 1 µm. *Right*: semi-automated quantification of Z-disk “disorder” as a fracture score (N=94–99 Z-disks analyzed; data as median ± 95% CI). **(E-K)** Transthoracic echocardiography (TTE) results in Het TC mice: (E) LV volumes (LVEDV and LVESV); (F) IVS dimensions (IVSd and IVSs); (G) LV posterior wall thickness (LVPWd and LVPWs); (H) Heart rate (HR) under anesthesia; (I) Stroke volume (SV); (J) Cardiac output (CO); (K) LV ejection fraction (LVEF). N=3 (Het TEV) and N=2 (Het GFP). Data are presented as mean ± SEM (except in panel D); statistical significance was determined by unpaired two-tailed t-test or Mann-Whitney test (panel D).

**Fig. S5.**
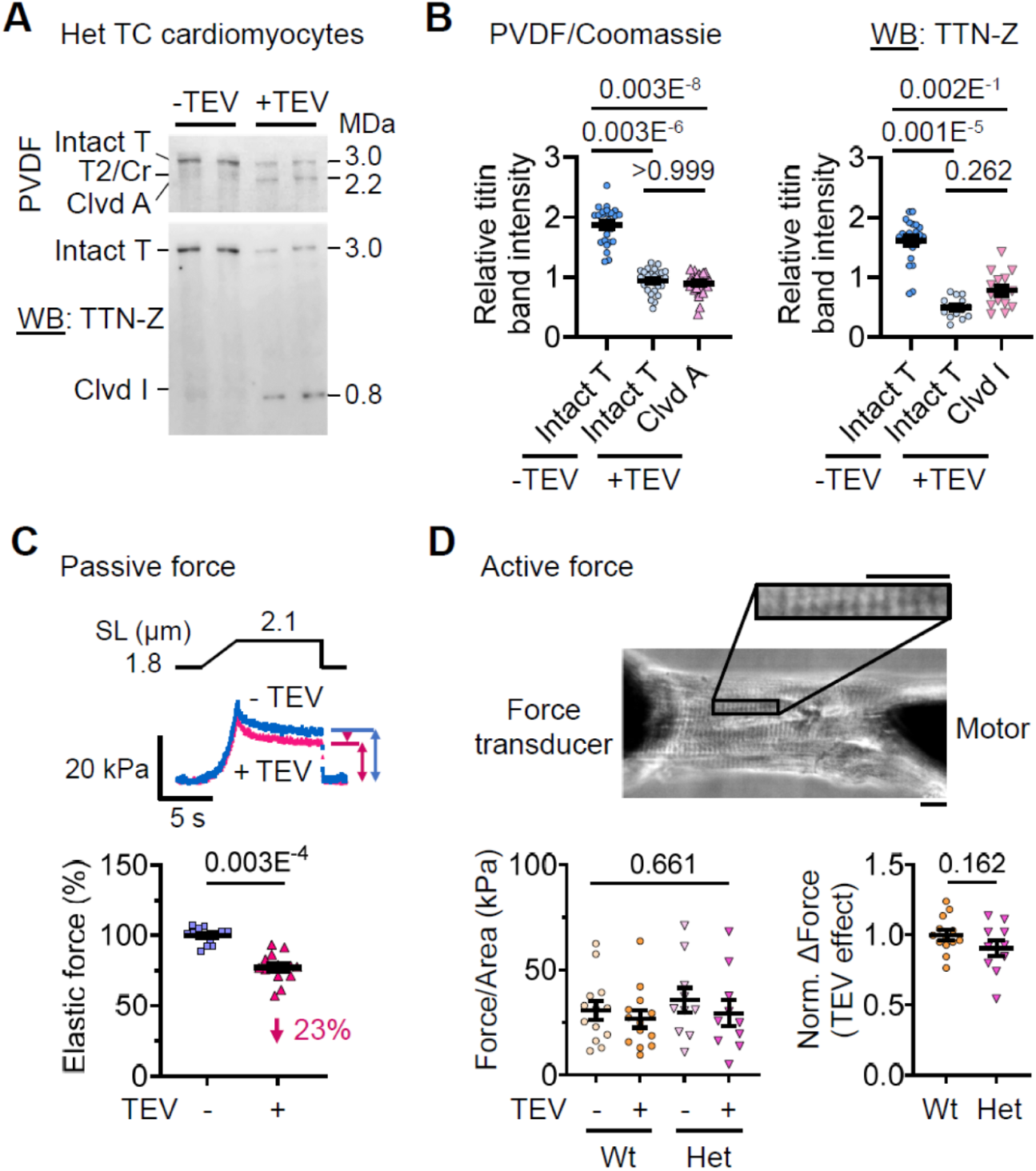
Effects of titin cleavage by TEV protease on passive and active forces of heterozygous (Het) TC mouse cardiomyocytes. **(A)** Titin protein gel (PVDF, *top*) and Western blot (WB, *bottom*) against Z-disk titin (TTN-Z) to quantify ex vivo titin cleavage by TEV in permeabilized Het TC left ventricular cardiomyocytes. **(B)** Densitometric analysis of titin band intensities showing ∼50% cleavage of titin after 30 min TEV incubation of permeabilized Het TC cardiomyocytes (n=28 for Coomassie, *left*; n=15 for WB, *right*). Statistical significance by Kruskal-Wallis test followed by Dunn’s multiple comparisons test. **(C)** Stretch protocol (*top*), representative passive force traces (*middle*), and average passive elastic force (*bottom*) of single, permeabilized Het TC cardiomyocytes before and after ex vivo TEV incubation, relative to pre-cleavage levels (n=15 cells/group); SL, sarcomere length. **(D)** Image of a permeabilized Het TC cardiomyocyte suspended between a force transducer and a micromotor (*top*), with a highlighted region of interest for SL analysis; scale bars, 10 µm. Graphs show average calcium-activated (pCa 5) force per cross-sectional area of single, permeabilized control wildtype (Wt) and Het TC cardiomyocytes before and after ex vivo TEV incubation (*left*) and average relative change of active force with TEV incubation of Het cardiomyocytes, compared to TEV effects on Wt control cells (*right*) (n=15 cells/group). All data are presented as mean ± SEM. Statistical significance was determined by unpaired two-tailed t-test or one-way ANOVA (D, *bottom*, *left*).

**Fig. S6.**
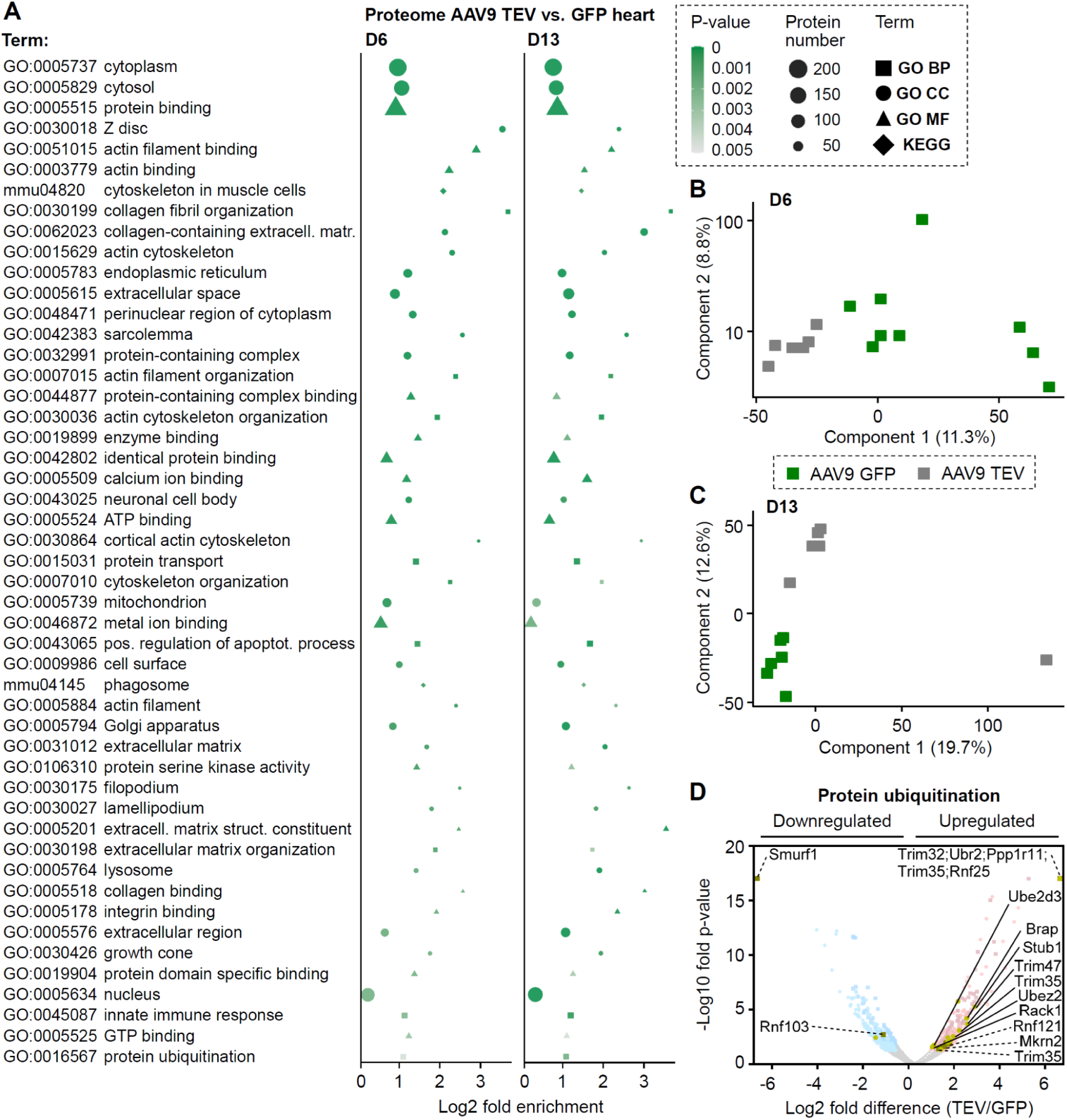
Additional proteomics analyses of titin-cleaved hearts. **(A)** GO and KEGG analyses of proteins significantly altered in TEV vs. GFP were performed using DAVID and visualized in R (ggplot2). Colors indicate a significance level of *p*=0.005, and shape size represents the number of proteins. GO terms are shown as BP (squares), CC (circles), MF (triangles); KEGG pathways as diamonds. Displayed terms, sorted by increasing *p*-value at D6, are those significant at both D6 and D13. **(B-C)** Principal component analysis of LV tissue proteomic profiles at (B) D6 and (C) D13 post-AAV9 injection. **(D)** Volcano plot of cardiac proteins in TEV vs. GFP at D6 (circles) and D13 (squares), showing proteins in the GO:0016567 (Protein ubiquitination) term (yellow); select proteins are identified via solid (D6) and dashed lines (D13). Faint background data refers to all proteins with a fold-change >1.5 (*p*<0.05), considered significantly up- or downregulated. D6: N=6 (Hom TEV), N=3 (Hom GFP), N=6 (Het GFP); D13: N=6 (Hom TEV and Hom GFP).

**Fig. S7.**
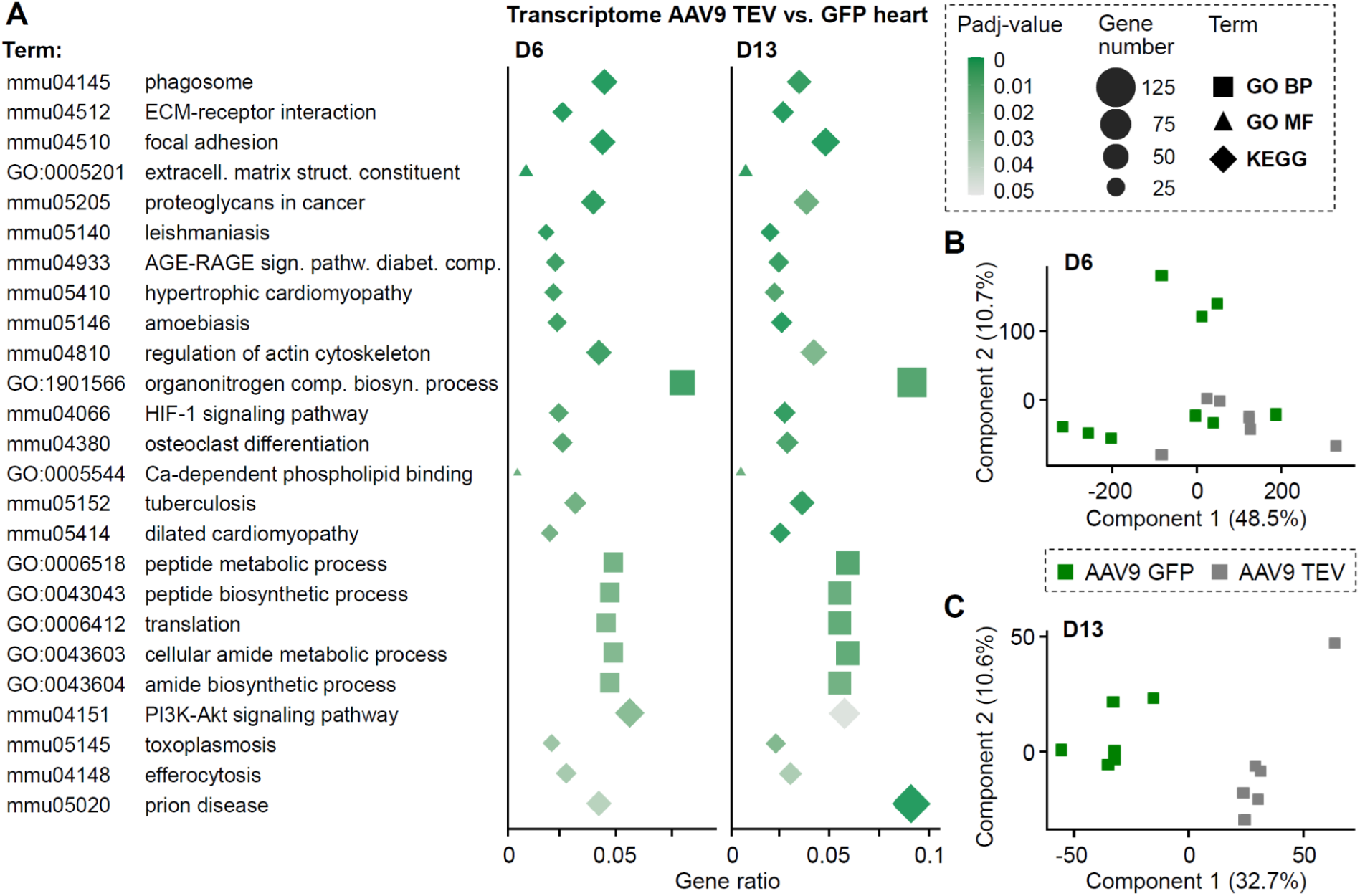
Additional transcriptomics of titin-cleaved mouse hearts. **(A)** GO and KEGG analyses of genes significantly altered in TEV vs. GFP were performed with DAVID and visualized in R (ggplot2). Color indicates *p*adj=0.05 significance and shape size represents gene count. GO BP is shown as squares, MF as triangles, and KEGG as diamonds. Displayed terms, ordered by increasing *p*-value at D6, are those significant at both D6 and D13. **(B-C)** Principal component analysis of LV transcriptomic profiles at (B) D6 and (C) D13 post-AAV9 injection. D6: N=6 (Hom TEV, Het GFP), N=3 (Hom GFP); D13: N=6 (Hom TEV and Hom GFP).

**Fig. S8.**
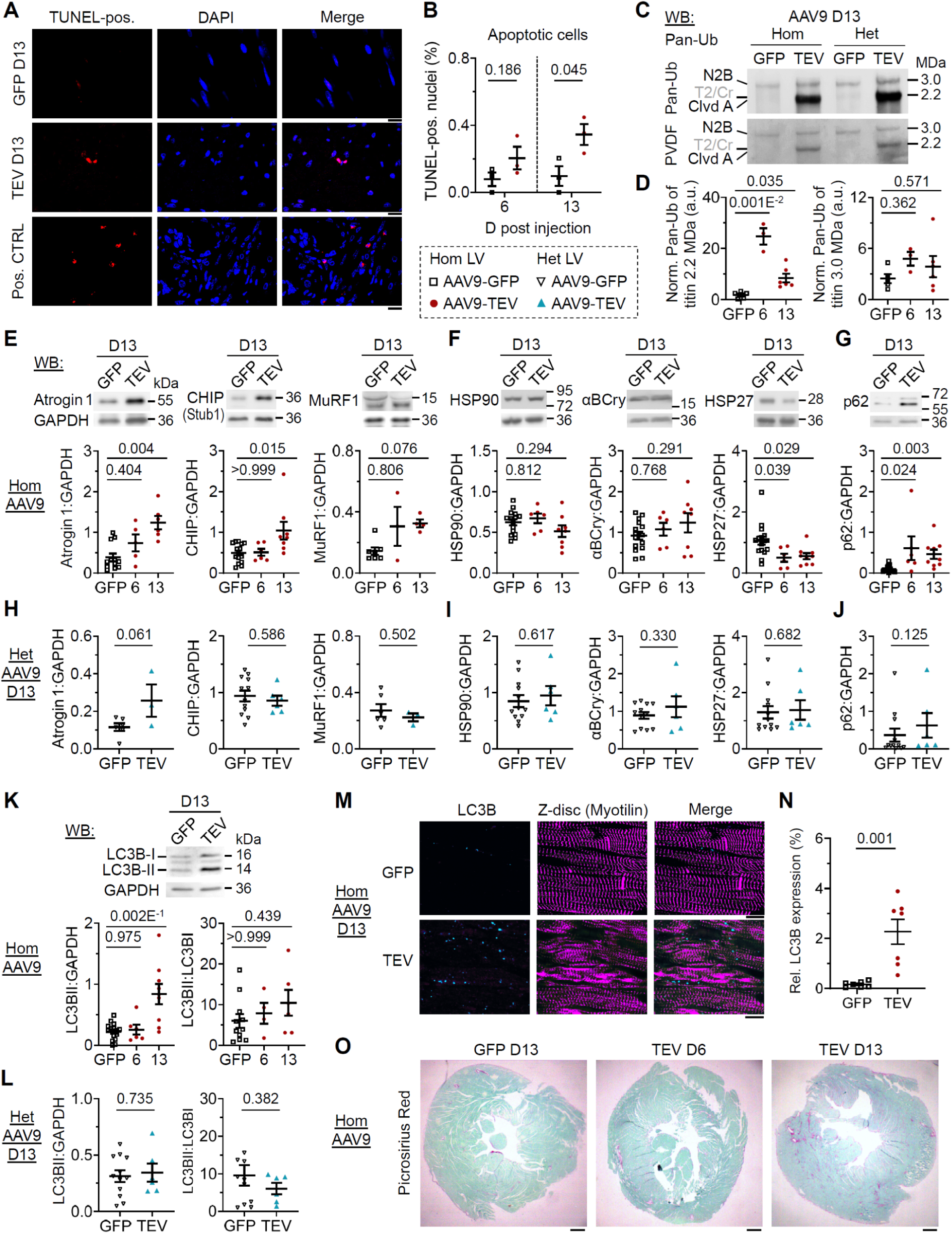
Markers of apoptosis, titin ubiquitination, protein quality control, and fibrosis in titin-cleaved heart tissue. **(A)** TUNEL staining of LV tissue from Hom TC mice at D6 and D13 post AAV9-GFP (*top*) or AAV9-TEV (*middle*) injection. Shown are TUNEL (*left*), DAPI (*middle*), and merged images (*right*). Mouse pancreatic tumor cells (*bottom*) served as a positive control. Scale bars, 20 µm. **(B)** Quantitation of TUNEL-positive cells as a percentage of DAPI-stained nuclei (N=3 mice/group). **(C)** Titin ubiquitination in AAV9-injected Hom (*left*) and Het (*right*) TC hearts at D13, assessed by anti-pan-ubiquitination Western blot (*top*) and Coomassie-stained PVDF (*bottom*). **(D)** WB quantification of titin pan-ubiquitination, normalized to Coomassie bands, for cleaved titin (2.2 MDa; *left*) and full-length N2B titin (3.0 MDa; *right*) in Hom TC hearts at D13 (N=5, GFP; N=6, TEV). **(E)** Expression of E3 ubiquitin ligases Atrogin1, CHIP (Stub1), and MuRF1 in Hom TC hearts, comparing AAV9-TEV mice at D6 and D13 to AAV9-GFP at D13 (representative WB; GAPDH control). **(F-G)** WB-based expression of chaperones (HSP90, αB-Crystallin, HSP27) (F) and autophagy adapter p62 (G) in Hom TC hearts. **(H-J)** WB-based expression of E3 ligases (H), chaperones (I), and p62 (J) in Het TC hearts at D13 (GFP vs. TEV). **(K-L)** WB-based expression of LC3BI/LC3BII in Hom (K) and Het (L) TC hearts, with summary data as LC3BII:GAPDH (*left*) and LC3BII:LC3BI (*right*) ratios (representative WB; GAPDH control). **(M)** Confocal images showing LC3B puncta (*left*) in Hom TC heart sections at D13 post AAV9-GFP (*top*) or AAV9-TEV (*bottom*) injection (Cy3), with Myotilin co-staining (*middle*; AlexaFluor 488) and merged images (*right*). Scale bars, 10 µm. **(N)** Relative LC3B puncta frequency per unit area in Hom TC hearts at D13 (n=7 images/group). **(O)** Picrosirius-red staining of whole heart sections from Hom TC mice at D6 and D13 post AAV9-TEV injection versus AAV9-GFP at D13. Scale bars, 0.5 mm. Data are shown as mean ± SEM. For WBs: N=8 (Hom GFP), N=5 (Hom TEV), N=6 (Het GFP), N=3 (Het TEV). Statistical significance was determined by unpaired, two-tailed t-test or Mann-Whitney test.

**Table S1.** Antibodies, dyes, and staining kits.

| Target antigen or structure | Vendor or Source | Catalog # | Working concentration |
| --- | --- | --- | --- |
| **Primary antibodies** |  |  |  |
| Anti-Alpha B Crystallin, mouse monoclonal | Abcam | Cat. #AB13496 | WB '(1:4000)' |
| Anti-CHIP/ STUB1, rabbit monoclonal | Abcam | Cat. #AB134064 | WB '(1:5000)' |
| Anti-Connexin 43, rabbit polyclonal | Abcam | Cat. #AB11370 | IF '(1:100)' |
| Anti-Cronos, rabbit polyclonal | Eurogentec | Custom-made | WB '(1:2000)' |
| Anti-Desmin D33, mouse monoclonal | Dako | Cat. #M 0760 | IF '(1:100)' |
| Anti-Desmoplakin, mouse monoclonal | Progen | Cat. #610035 | IF '(1:5)' |
| Anti-FBXO32 (Atrogin1), rabbit polyclonal | St John's Laboratory | Cat. #STJ23637 | WB '(1:1000)' |
| Anti-GAPDH (14C10), rabbit monoclonal | Cell Signaling | Cat. #2118 | WB '(1:40000)' |
| Anti-GFP, monoclonal rabbit | Cell Signaling | Cat. #2956 | WB '(1:1000)' |
| Anti-HSP27, rabbit monoclonal | Developmental Studies Hybridoma Bank | Cat. #CPTC-HSPB 1-4 | WB '(1:1000)' |
| Anti-HSP90, rabbit polyclonal | Cell Signaling | Cat. #4874 | WB '(1:1000)' |
| Anti-Ki67, rabbit polyclonal | Abcam | Cat. #ab15580 | IF '(1:250)' |
| Anti-LC3B, rabbit polyclonal | Cell Signaling | Cat. #2775 | IF '(1:200)'<br>WB '(1:2500)' |
| Anti-MuRF1 (C-11), mouse monoclonal | Santa Cruz | Cat. #sc-398608 | WB '(1:10000)' |
| Anti-Myotilin (Leica NLC Myotilin), mouse monoclonal | ThermoFisher Scientific | Cat. #50-255-2199 | IF '(1:100)' |
| Anti-N-Cadherin, mouse monoclonal | Invitrogen | Cat. #33-3900 | IF '(1:100)' |
| Anti-p62, rabbit monoclonal | Abcam | Cat. #AB109012 | WB '(1:2500)' |
| Anti-Pan-Ubiquitin P4D1, mouse monoclonal | Santa Cruz | Cat. #sc-8017 | WB '(1:100)' |
| Anti-PCM1, rabbit polyclonal | Merck Millipore | Cat. #HPA 023374 | IF '(1:250)' |
| Anti-Periostin, mouse monoclonal | Proteintech | Cat. #66491-I-Ig | IF '(1:1000)' |
| Anti-TEV, rabbit polyclonal | Abcepta | Cat. #ABV11239-100 | WB '(1:500)' |
| Anti-TEV cleavage site (cTEV), rabbit polyclonal | ThermoFisher Scientific | Cat. #PA1-119 | IF '(1:500)'<br>WB '(1:1000)' |
| Anti-Titin I20-22, rabbit polyclonal | Myomedix | Cat. #TTN-5 | IF '(1:400)'<br>WB '(1:2000)' |
| Anti-Titin M, rabbit polyclonal | Myomedix | Cat. #TTN-9 | IF '(1:100)' |
| Anti-Titin MIR, rabbit polyclonal | Myomedix | Cat. #TTN-6 | WB '(1:20000)' |
| Anti-Titin Z, rabbit polyclonal | Myomedix | Cat. #TTN-1 | WB '(1:10000)' |
| **Secondary antibodies** |  |  |  |
| AlexaFluor 488 affininpure goat anti-mouse IgG, goat polyclonal | Jackson ImmunoResearch | Cat. #115-546-146 | IF '(1:250)' |
| Cy3 affininpure goat anti-rabbit IgG, goat polyclonal | Jackson ImmunoResearch | Cat. #111-165-003 | IF '(1:250)' |
| **Dyes and staining kits** |  |  |  |
| DAPI | ThermoFisher Scientific | Cat. #D1306 | '(1:500)' |
| OneStep TUNEL Apoptosis Kit (Red, 555) | Novus Biologicals | Cat. #NBP3-12093 |  |
| Sirius Red/Fast Green collagen staining kit | Biozol | Cat. #9046 |  |
| Wheat germ agglutinin, AlexaFluor 555 conjugated | Invitrogen | Cat. #W32464 | '(1:250)' |
| Wheat germ agglutinin, AlexaFluor 647 conjugated | Invitrogen | Cat. #W32466 | '(1:500)' |

**Movie S1.**

This movie shows long-axis cardiac MRI frames recorded from a TC mouse injected with AAV9-GFP on D6 post injection.

**Movie S2.**

This movie shows long-axis cardiac MRI frames recorded from a TC mouse injected with AAV9-TEV on D6 post injection.

**Movie S3.**

This movie shows long-axis cardiac MRI frames recorded from a TC mouse injected with AAV9-TEV on D13 post injection.

**Data S1-S5 (separate file)**

**Data S6-S13 (separate file)**

**Data S14 (separate file)**

